# Ciliated cells promote high infectious potential of influenza A virus through the efficient intracellular activation of hemagglutinin

**DOI:** 10.1101/2025.04.30.651529

**Authors:** Zijian Guo, Victoria S. Banas, Yuanyuan He, Elizabeth Weiland, Jian Xu, Yangjie Tan, Zhaoxi Xiao, Steven L. Brody, Adrianus C. M. Boon, James W. Janetka, Michael D. Vahey

## Abstract

Influenza viruses utilize host proteases to activate the viral fusion protein, hemagglutinin (HA), into its fusion-competent form. Although proteolytic activation of HA is essential for virus replication, the cell-type dependence of HA activation within the airway epithelium and the subcellular location(s) in which it occurs are not well-established. To address these questions, we investigated the proteolytic activation of HA in differentiated human airway epithelial cells using contemporary and historical H1N1 and H3N2 strains. We find that activation is efficient across viral strains and subtypes but depends on cellular tropism, with ciliated cells activating HA more effectively than non-ciliated cells. Similar to prior observations in immortalized cell lines, we find that HA activation occurs intracellularly, constraining the antiviral activity of host-directed protease inhibitors. These results establish that HA activation within the airway epithelium depends on cellular tropism, and identify important considerations for the development of protease inhibitors as antivirals.

**Importance:** Influenza entry requires the proteolytic activation of the viral fusion protein, hemagglutinin (HA). Activation occurs as new viruses are produced by infected cells. Efficient proteolytic activation is critical for viral pathogenesis, and inhibiting the requisite proteases could provide an effective host-directed antiviral strategy. To understand cellular constraints on HA activation and its sensitivity to inhibitors, we use complementary approaches to investigate these processes in differentiated airway epithelial cells. We find that ciliated cells activate HA with higher efficiency than non-ciliated cell types, establishing a new mechanism through which cellular tropism and virus infectious potential are connected. We also establish that HA activation begins in the Golgi, which may contribute to the limited the efficacy of inhibitors we observe despite their high *in vitro* potency.

## Introduction

Enveloped viruses use fusion proteins on the viral surface to deliver their genomes across cellular membranes. For enveloped viruses with class I and class II fusion proteins, proteolytic processing of the fusion protein by host proteases is a prerequisite for cellular entry^1^. This processing step enables the extensive refolding of the fusion protein in response to environmental cues, bringing the viral and host membrane in close apposition and lowering the kinetic barrier to membrane fusion^2^. Respiratory viruses including influenza, coronaviruses, and pneumoviruses have evolved to utilize an overlapping repertoire of host proteases^3^. While viruses possessing polybasic motifs at their cleavage sites can utilize the ubiquitous serine protease furin^4–7^, monobasic cleavage sites are processed by alternate serine proteases. Protease dependency is thought to be an important determinant of virulence and tissue tropism, constraining the cell types in which a virus can replicate^8,9^.

The fusion protein of influenza viruses, hemagglutinin (HA), can possess either a polybasic or monobasic cleavage site. While polybasic motifs are associated with highly pathogenic viruses, seasonal influenza A viruses currently circulating in humans contain monobasic cleavage sites^10^. Although the activation of HAs with monobasic cleavage sites can be achieved by a wide variety of serine proteases including exogenous trypsin^11–13^, multiple lines of evidence suggest that TMPRSS2 is the major activating protease in humans and mice. Knockdown of TMPRSS2 using an antisense morpholino prevented HA activation and replication of both H1N1 and H3N2 strains in primary human bronchial epithelial cells^14^. While additional studies in TMPRSS2 knockout mice corroborate these findings^15^, the protease dependencies of H3N2 strains in mice are less well defined^16,17^.

The fusion proteins of different viruses are proteolytically activated at different stages of the replication cycle. While the processing of SARS-like coronavirus spike proteins occurs in *trans* during viral entry by surface-associated or endosome-associated proteases of the target cells^18,19^, influenza HA is processed by proteases expressed by the infected cell or present in the extracellular environment^20^. This distinction carries important implications in the human airways, where the epithelium is comprised of a complex mosaic of different cell types which differ both in their susceptibility to infection as well as their ability to produce infectious virus^21,22^. Accordingly, the ability of distinct airway cell populations to support proteolytic activation of HA may contribute to cross-species barriers to transmission^23^. Moreover, prior studies have reached different conclusions regarding whether HA activation predominantly occurs intracellularly, at the cell surface, or within the extracellular space^24,25^. Identifying the site of HA activation in differentiated airway epithelial cells could guide the development of antiviral protease inhibitors that are able to reach the subcellular locations where activation occurs.

To better define the cellular and sub-cellular landscape of proteolytic activation of influenza HA in the human airways, we used a combination of biochemical and imaging-based approaches to map proteolytic activation of H1N1 and H3N2 strains within differentiated human airway epithelial cells. We find that proteolytic activation of HA is efficient across strains, and that virus infectious potential is highly sensitive to modest reductions in activation efficiency. We also demonstrate that HA activation in differentiated airway epithelial cells begins in the Golgi, and that a range of virus- and host-directed inhibitors of HA activation differ markedly in their potency – likely due to differences in their ability to access intracellular pools of activating proteases. Finally, we show that HA activation differs between cell types within the airway epithelium, with multi-ciliated cells showing the most efficient activation. Thus, proteolytic activation of HA provides a means through which the infectious potential of viruses produced in the airway epithelium is inherently linked to cellular tropism.

## Results

### Differentiated airway epithelial cells efficiently activate H1 and H3 HAs

To determine the efficiency of HA activation in a model human airway epithelium, we differentiated primary human tracheal epithelial cells (HTECs) at the air-liquid interface (ALI) for >4 weeks. These cultures differentiate into a complex epithelium that models the human airways *in vivo*^26^. After removing mucus by apical wash and treatment with the enzyme StcE^27^, we infected HTEC cultures with H1N1 (A/California/04/2009 or A/WSN/1933) or H3N2 (A/Brisbane/10/2007 or A/Hong Kong/1968) strains at a multiplicity of infection (MOI) of ∼0.5 for 1 h before washing away the inoculum. At 20 h post-infection (h.p.i.), we collected released viruses via apical wash and quantified the extent of HA activation via Western blot (Figure 1A & Supplementary Figure 1). To extract a numerical value for HA activation from the blots, we calibrated the results with a standard sample that is composed of an equal mixture of HA0 and fully-activated HA (Supplementary Figure 2). This allows us to control for differences in the transfer and detection of HA0 and HA1/HA2 during Western blotting. Quantification of HA activation revealed that shed virus from each strain was efficiently activated (>90%) (Figure 1B).

**Figure 1:**
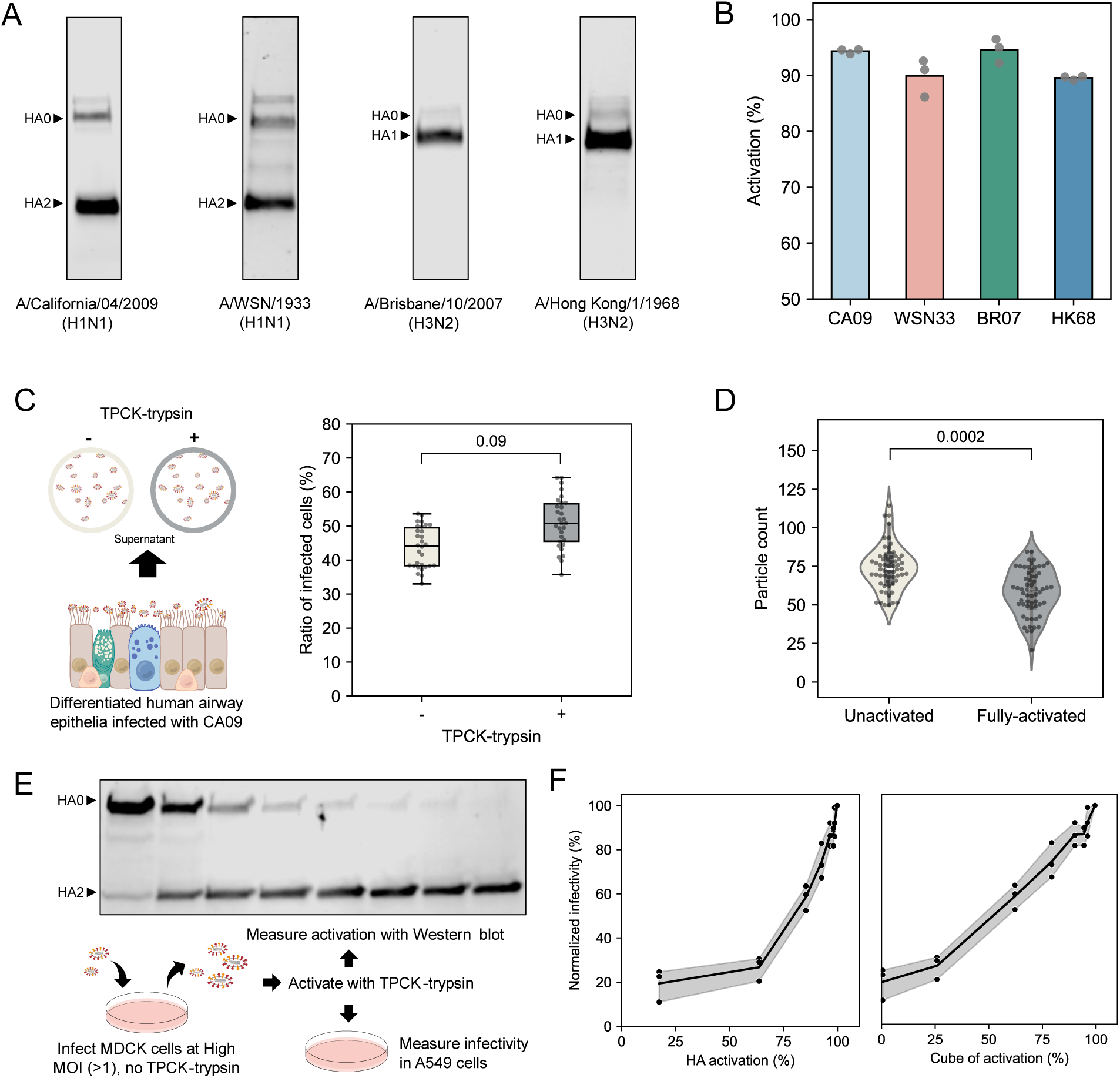
Efficient activation of HA in airway epithelial cells supports high infectious potential. (A) Representative images of Western blots of H1 and H3 HAs. (B) Quantification of activation of H1 and H3 strains. Data represent three infection experiments performed in separate cultures from the same donor. (C) Schematic and quantification of infectivity for virus collected from differentiated HTECs +/− further activation by TPCK-treated trypsin. Data are combined from virus samples grown in three HTEC cultures derived from a single donor. Each data point represents one field of view of the fluorescence confocal image. The p-value is determined by an independent t-test on the mean values of all fields of view in each independent experiment. (D) Quantification of binding of unactivated and fully-activated viruses grown from MDCK cells. Experiments were conducted using the same virus samples, with six A549 cultures used as target cells for virus binding in each condition. Each data point represents one field of view of the fluorescence confocal image. The p-value is determined by an independent t-test on the mean values of all fields of view in each independent experiment. (E) Western blot and schematic showing the production and testing of viral stocks with variable activation levels. (F) Viral infectivity as a function of percentage HA activation for CA09. Left plot: HA activation displayed as percentage of activated monomers. Right plot: HA activation displayed as the predicted ratio of fully-activated HA trimers, identified by the cube of HA activation. Data represent three independent experiments with virus diluted from the activation series shown in E.

Although only ∼5% of viral HA from HTEC cultures infected with A/California/04/2009 (‘CA09’) remained unactivated, subsequent treatment of this virus with TPCK-trypsin increased infectious potential by an average of ∼23% (Figure 1C). The disproportionate change in infectivity is not due to changes in virus binding, which is similar between fully-activated and unactivated viruses (Figure 1D). To determine more quantitatively the relationship between HA activation and viral infectivity, we prepared unactivated CA09 virus from MDCK cells and titrated viral activation by incubating with TPCK-trypsin for different durations before adding a five-fold molar excess of soybean trypsin inhibitor and measuring virus infectious potential on A549 cells (Figure 1E). Consistent with our findings for virus produced by differentiated HTECs, we observe disproportionate increases in infectious potential when the percentage of activated HA monomers exceeds ∼60% (Figure 1F & Supplementary Figure 3). We reasoned that unactivated HA may restrict infection by disrupting the coordinated extension or refolding of HA trimers that contain one or more unactivated monomer(s) during membrane fusion^28^. Assuming that activated HA monomers are randomly distributed among trimers, we expect the fraction of HA trimers in which all monomers are cleaved to scale as the cube of the fraction of activated HA monomers. Rescaling our data in this way reveals a roughly linear relationship between infection and trimer activation. This sensitivity to unactivated HA suggests that even modest inhibition could strongly reduce viral replication in the airway epithelium.

### Virus- and host-directed strategies to inhibit HA proteolytic activation show variable potency

The HA cleavage site is located membrane-proximally, where it occupies a cleft between adjacent monomers within the trimer and partly occludes the anchor epitope^29,30^. Antibodies that bind to membrane-proximal regions of HA may be capable of preventing proteolytic activation^31,32^. Using AlphaFold2^33^, we modeled full-length trimeric HA0 (from CA09) in complex with the processed TMPRSS2 extracellular domain and transmembrane domain. As expected, this model positioned the P1 Arg from HA1 within the S1 pocket of TMPRSS2, and positioned the P1’ Gly from HA2 where it is accessible for nucleophilic attack from the catalytic serine (Figure 2A). The transmembrane domains of HA and TMPRSS2 are also aligned, providing a plausible model for how they might sit in the membrane during activation. Aligning this model to structures of the stalk- and anchor-binding antibodies FI6v3 and FISW84 suggests that binding of either antibody would sterically occlude recognition of the cleavage loop by TMPRSS2 and likely other serine proteases (Figure 2B & Supplementary Figure 4). We tested the effects of these antibodies on HA activation using unactivated CA09 virus produced in MDCK cells. While both FISW84 Fab and IgG showed partial inhibition of HA activation when they were pre-incubated with virus before adding TPCK-trypsin, we did not observe inhibition by FI6v3 under the conditions tested, although it was previously verified to block activation of recombinant HA^32^ (Figure 2C Left). We proceeded to test the effects of FISW84 IgG in differentiated HTECs, delivered both apically and basally at a concentration of 1 µM following infection. Although this concentration is ∼40-fold above the IC_50_ of FISW84 for viral entry^30^, we did not observe noticeable changes in HA activation (Figure 2C Right). This suggests that the accessibility of HA prior to activation may limit the effectiveness of HA-specific antibodies in restricting proteolytic processing.

**Figure 2:**
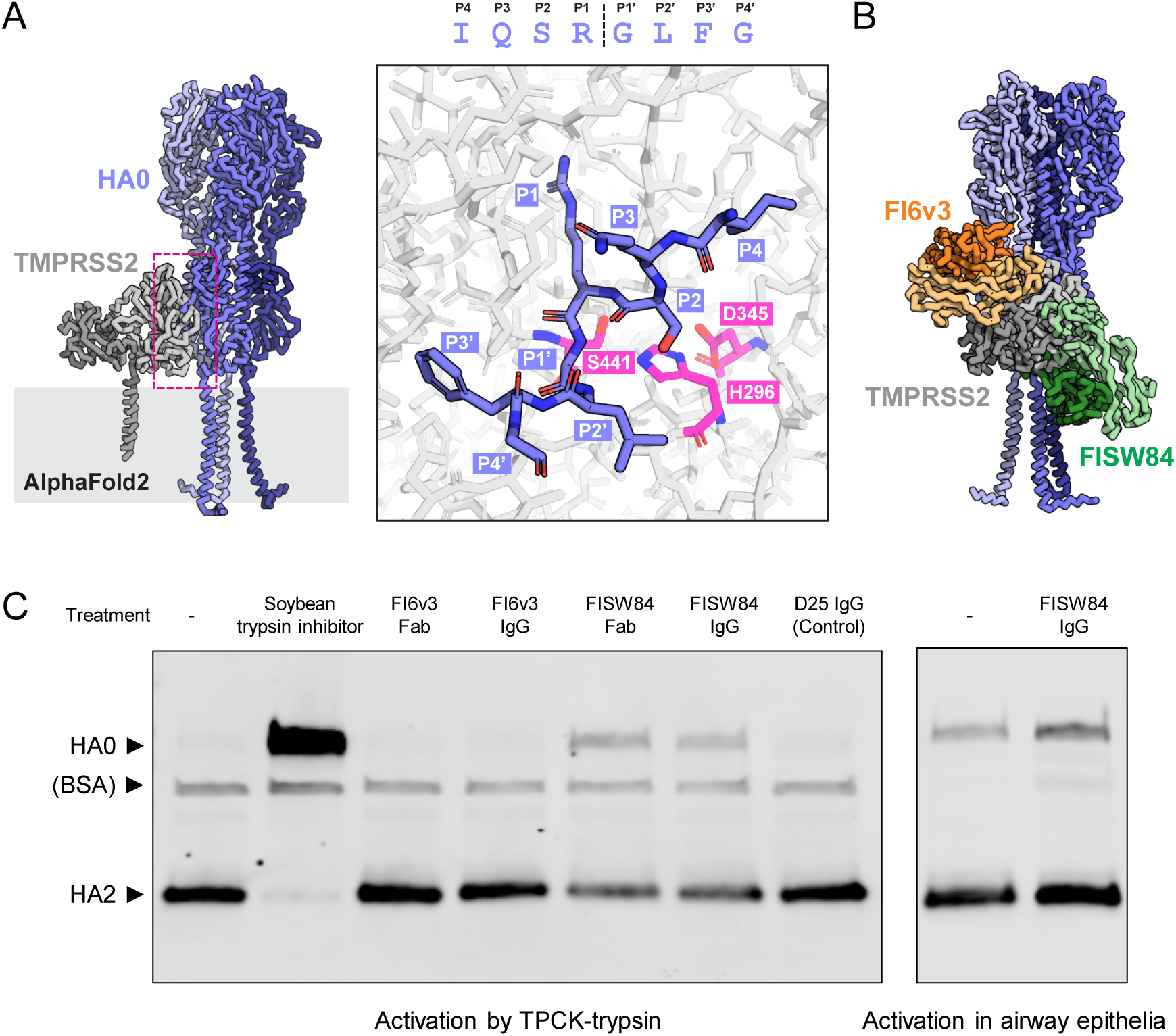
Monoclonal antibodies that occlude the HA cleavage site are inefficient inhibitors of HA activation in airway epithelial cells. (A) Left: AlphaFold2 model depicting H1 HA0 in complex with TMPRSS2. Right: en face view of the HA cleavage loop (blue) positioned within the active site of TMPRSS2 (catalytic residues shown in magenta). (B) Model of stem-binding FI6v3 and anchor-binding FISW84 Fab fragments superimposed on the structural prediction from A. (C) Western blot showing inhibition of HA activation by FI6v3 and FISW84 antibodies and Fab fragments. Blot to the left is for unactivated virus produced in MDCK cells and subsequently treated with TPCK-trypsin, +/− the indicated treatments. Blot to the right shows representative results from differentiated HTEC cultures.

Direct inhibition of host proteases^34^ provides an alternative to HA-directed approaches to prevent the proteolytic activation of influenza HA. We compared the efficacy of a panel of protease inhibitors in differentiated HTEC cultures by delivering them both apically and basally following infection and collecting virus at 20 h.p.i. for analysis (Figure 3 & Supplementary Figure 5). To attempt to isolate the intrinsic potency of each inhibitor from its ability to access its target protease(s) in differentiated HTECs, we tested each inhibitor at concentrations >10-fold above the *in vitro* IC_50_ value, as determined against recombinant TMPRSS2 extracellular domain (Supplementary Figure 6 & Supplementary Figure 7). Among the candidates we tested, Nafamostat, a cell permeable broad-spectrum serine protease inhibitor^35^, showed complete inhibition of H1 activation and strong inhibition of H3 activation at 100 nM. In comparison, MM3122, a peptidomimetic drug with subnanomolar potency against TMPRSS2 and matriptase^36^ showed strong but incomplete inhibition of H1 activation, but had little effect on activation of H3 HA. We also tested three protein-based inhibitors: two Kunitz-type inhibitors (Soybean trypsin inhibitor^37^, STI, and Hepatocyte growth factor activator inhibitor-1^38,39^, HAI-1), as well as a TMPRSS2 nanobody^40^ (A07). Although each of these inhibitors exhibited potency against soluble TMPRSS2, they showed only modest inhibition of HA activation in differentiated HTECs for both viral subtypes. In a separate experiment, we also tested inhibition by plasminogen activator inhibitor (PAI-1); although recombinant PAI-1 protected against HA activation by exogenous trypsin, it did not prevent HA activation in differentiated HTECs (Supplementary Figure 8). Recombinant PAI-1 has previously been shown to inhibit extracellular activation of WSN33^25^, suggesting its inhibitory effects in this form may be strain-dependent. Similarly, the reduced susceptibility of BR07 HA activation to protease inhibitors with higher specificity towards TMPRSS2 (MM3122, A07) aligns with the subtype-specific protease dependencies observed by others^16,41^. Differences in the relative potency of these inhibitors in HTECs versus in vitro may result from differences in their membrane permeability, which could limit access to the pool of proteases responsible for HA activation.

**Figure 3:**
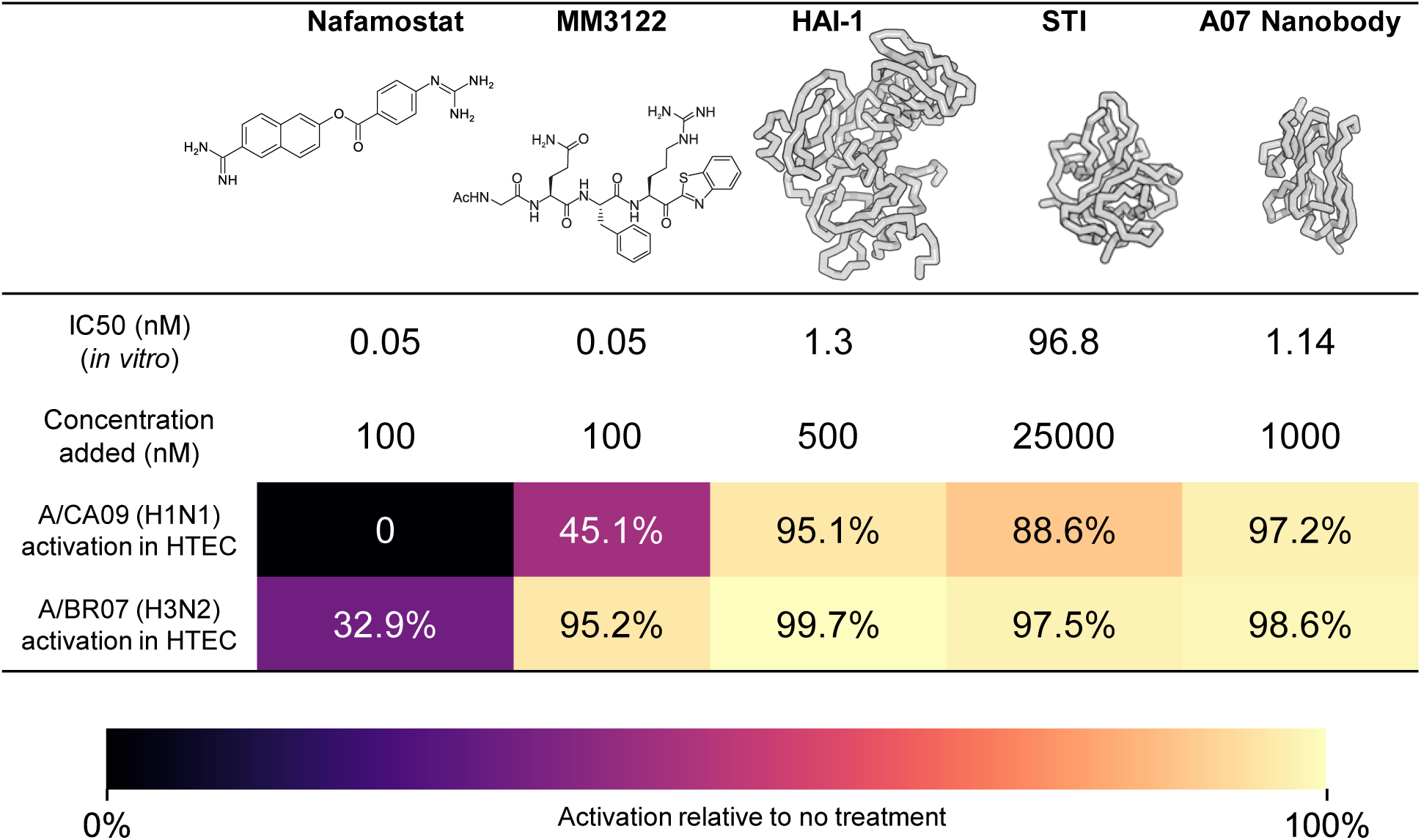
Inhibition of HA activation by Nafamostat, MM3122, HAI-1, Soybean trypsin inhibitor (STI) and A07 anti-TMPRSS2 nanobody. IC_50_ values are measured against TMPRSS2 ectodomain *in vitro* and show the average of two independent trials. Inhibition data are from three independent experiments performed with differentiated HTECs from different donors. Virus from a non-treated culture for each donor is used as negative control.

### HA activation begins early in the secretory pathway

Interestingly, when we increased the concentration of STI further to 250 μM (>2500 times its IC_50_ against recombinant TMPRSS2), we observed more pronounced inhibition (Figure 4A). We hypothesized that this enhanced inhibition results from improved targeting of a pool of proteases that recycle from the plasma membrane to endosomal compartments. This hypothesis is supported by live-cell imaging of airway cultures, which reveal intracellular localization of fluorescently labeled STI after 45 minutes of incubation, in addition to its presence on the cell surface and the ciliary membrane (Figure 4B). Altogether, these results suggest that HA may undergo intracellular activation prior to reaching the cell surface or assembling into budding virions. Consequently, membrane-impermeable inhibitors would require high concentrations or long exposure to effectively block their target proteases, a potential limitation in their therapeutic application.

**Figure 4:**
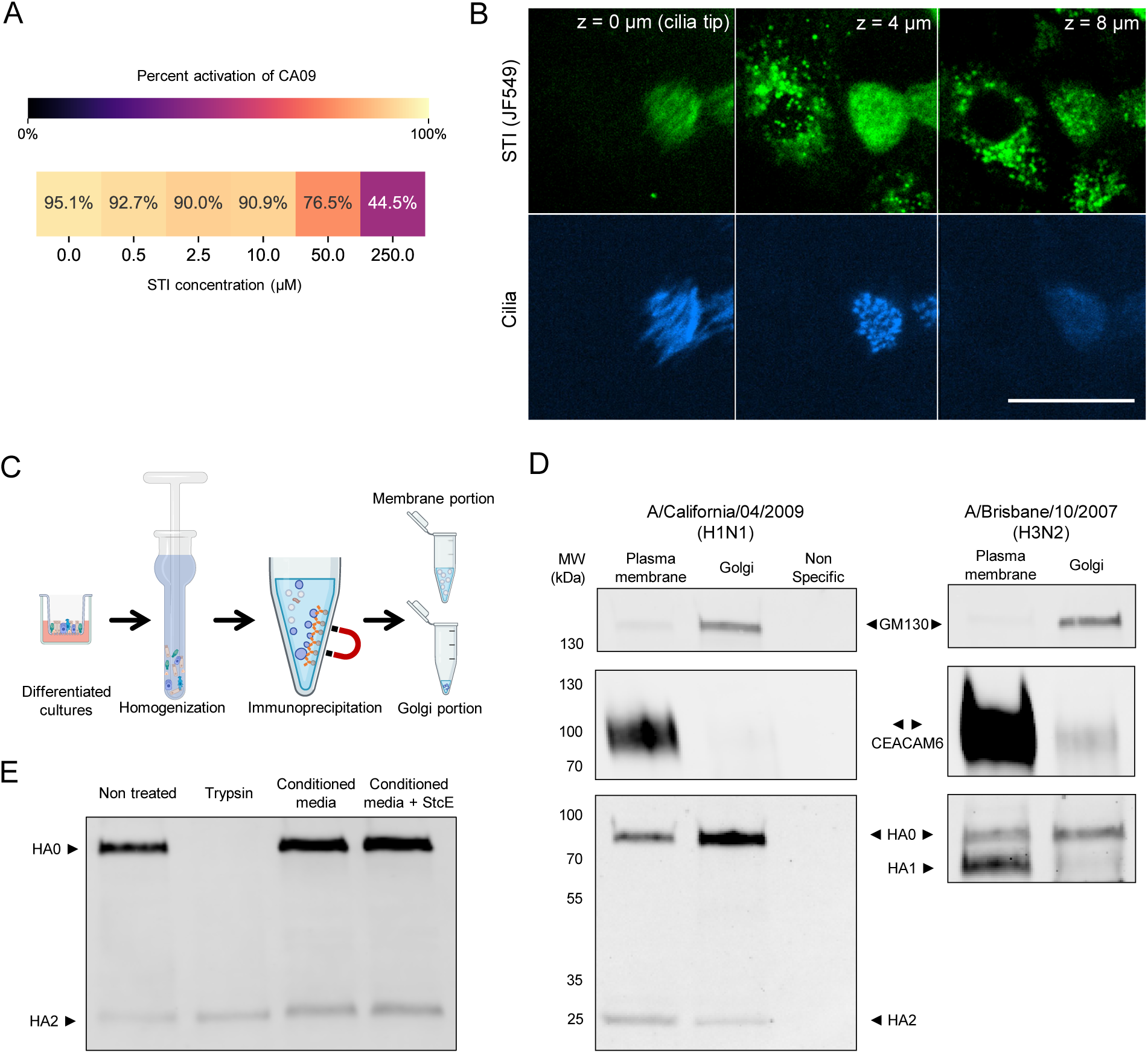
HA activation begins in the Golgi of differentiated airway epithelial cells. (A) Concentration-dependent inhibition of HA activation in airway cultures by soybean trypsin inhibitor. (B) Live imaging of differentiated HTECs incubated with fluorescent soybean trypsin inhibitor (JF549) and SiR-Tubulin. Images depict the same x-y position at different heights (z) relative to the cilia. Scale bar = 25 μm. (C) Schematic illustrating the isolation of native Golgi membranes from differentiated HTECs. (D) Western blot of immunoprecipitated Golgi membranes from differentiated HTEC cultures infected with CA09 (left) or BR07 (right). Non-specific lane corresponds to beads loaded with anti-RSV IgG. CEACAM6 is used as a marker of the non-Golgi membrane fraction. (E) Western blot of initially unactivated virus from MDCK cells following control treatment, treatment with trypsin, or treatment with apical secretions from differentiated cultures (+/− the mucinase StcE). Experiment is performed with conditioned media from cells of donor WU262. Data from another donor is shown in Supplementary Figure 10.

Prior studies using MDCK cells expressing human airway proteases^24^ and Caco-2 human intestinal cells^42^ demonstrate that HA activation can occur during the secretory pathway. To determine if this is the case in differentiated HTECs, we immunoprecipitated native Golgi membranes from differentiated cultures following infection with CA09 and BR07 and evaluated the activation status of HA via Western blot (Figure 4C). For both strains, we detect proteolytic processing of HA within membranes immunoprecipitated by the cis-Golgi marker GM130^43^, although to a lesser extent than the crude membrane fraction. Consistent with this observation, immunofluorescence of TMPRSS2 and GM130 shows partial colocalization in both infected and uninfected cultures, with HA visible in the Golgi of infected cells (Supplementary Figure 9). These results suggest that HA activation begins as early as the cis-Golgi compartment and continues into later stages of the secretory pathway.

Serine proteases are cleaved during zymogen activation and may undergo additional cleavage resulting in shedding of the serine protease domain^44–46^. To determine if secreted proteases are likely to contribute to HA activation in our experiments, we collected apical washes from differentiated cultures (20 µL of Hank’s buffered salt solution, HBSS, per insert, incubated for 1 h at 37°C) and mixed these 1:1 with unactivated CA09 virus for 12 h at 37°C before analyzing HA activation by Western blot. We found similar levels of HA activation regardless of whether virus was incubated with HBSS or HBSS plus apical secretions (Figure 4D & Supplementary Figure 10). This result was not affected by the presence or absence of StcE^27^ which helps extract virions from mucus, suggesting that the observable mucus gel collected with the wash did not protect virus from activation. These results suggest that, in the absence of secreted protease activity, HA activation falls within the jurisdiction of the infected cell and is constrained by the collection of proteases it expresses.

### HA activation depends on the differentiation status of airway epithelial cultures

The human airway epithelium is comprised of multiple cell types that express distinct repertoires of proteases. Using the Human Cell Atlas consortium respiratory single cell RNA-seq data^47^, we compared the expression of proteases previously identified to activate influenza HA: TMPRSS2, TMPRSS4, matriptase/ST14, HAT/TMPRSS11D, and furin (Figure 5A). Goblet and club cells show the highest expression of most proteases, with TMPRSS4 most highly expressed across cell types, as previously observed^14^. Heterogeneity in protease expression is also captured by immunofluorescence staining of TMPRSS2 in differentiated HTECs, which shows localization in the cilia and apical membrane of ciliated cells and intracellular and basolateral membranes in non-ciliated cells (Figure 5B).

**Figure 5:**
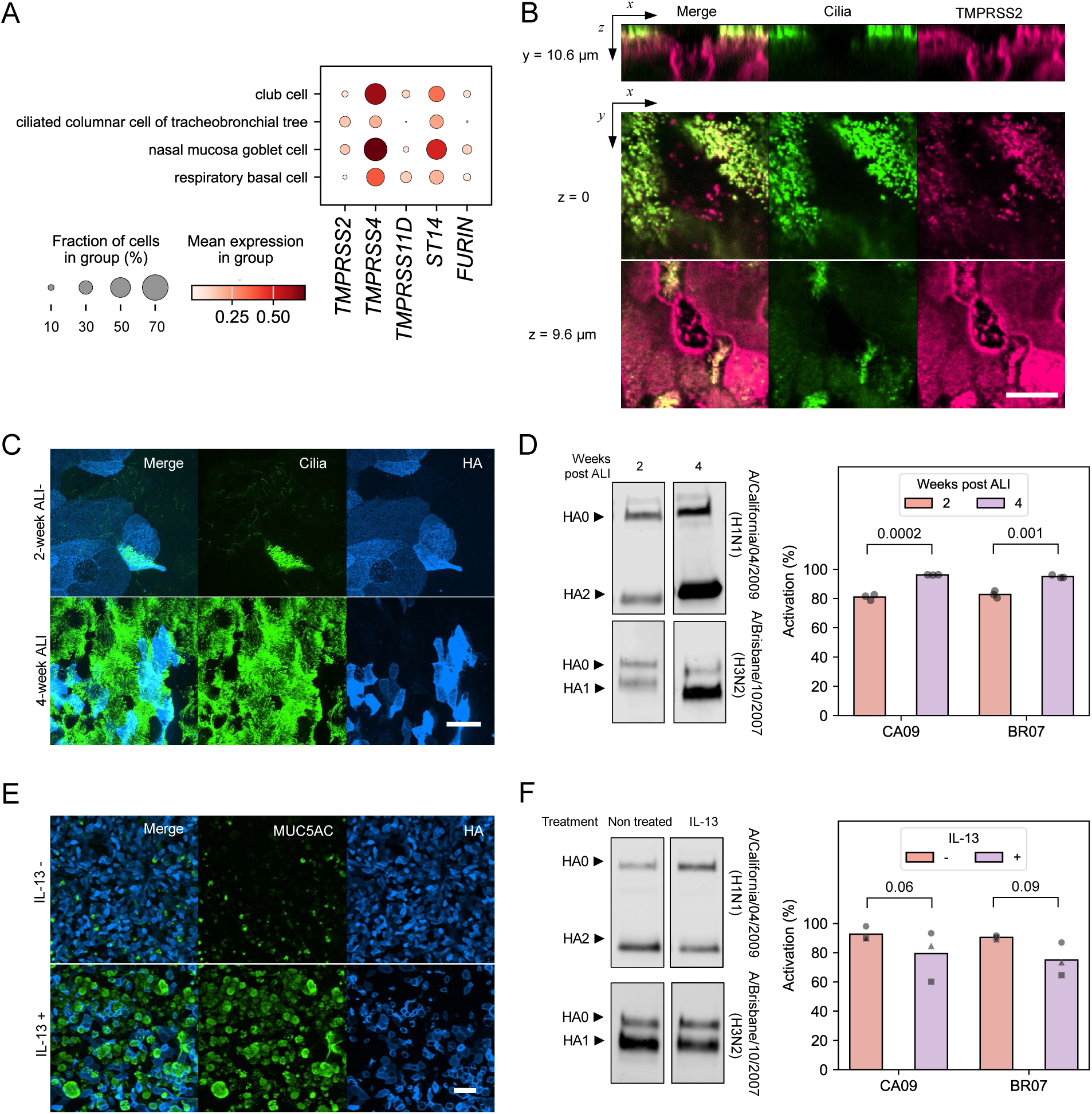
HA activation correlates with ciliation. (A) Expression analysis of TMPRSS2, TMPRSS4, TMPRSS11D, ST14 (Matriptase) and FURIN in different cell types within airway epithelia from the Human Lung Cell Atlas^64^. (B) Immunofluorescence confocal images showing the expression of TMPRSS2 in ciliated and non-ciliated cells. The thickness of the *xz* view is 20.7 μm. Scale bar = 10 μm. (C) Immunofluorescence confocal images showing differences in ciliation for 2-week and 4-week post-ALI samples. Cells were infected with BR07 (H3N2) and labeled with FI6v3 anti-HA Fab at 20 h.p.i. Scale bar = 25 μm. (D) Western blot and quantification of HA activation in CA09 and BR07 viruses produced from 2-week and 4-week post-ALI cultures. Data are from three independent infections of separate cultures from one donor performed on the same day. P-values are determined by independent t-tests. (E) Immunofluorescence confocal images showing differences in MUC5AC expression in untreated and IL-13-treated HTECs. Cells were infected with BR07 (H3N2). Scale bar = 50 μm. (F) Western blot and quantification of HA activation in CA09 and BR07 viruses produced from HTEC cultures +/− IL-13 treatment to modulate MUC5AC+ cell differentiation. Data are from three experiments, where a culture of IL-13 treated and another untreated were infected side-by-side. P-values are determined by paired t-tests on samples infected at the same time.

We reasoned that differences in protease expression and localization could affect the efficiency of HA activation in different cell types. As an initial test, we compared activation in undifferentiated (2 weeks after ALI) and well-differentiated (4 weeks after ALI) cultures showing different degrees of ciliation (Figure 5C). Viruses from well-differentiated cultures exhibited higher HA activation (Figure 5D). To further investigate cell-type dependent activation, we produced HTEC cultures with skewed differentiation. Our standard differentiation conditions produce cells with high degrees (>80%) of ciliation at 4 weeks post-ALI. Treatment with IL-13 shifts differentiation towards a mucous (MUC5AC+) cell phenotype^48,49^ (Figure 5E). For both CA09 and BR07 strains, we observe reduced HA activation in virus shed by IL-13-treated cultures, although the result does not reach statistical significance (Figure 5F). Collectively, these results suggest that the degree of HA activation correlates with ciliation.

### An imaging-based assay reveals ciliated cells as the most efficient activators of HA

To examine cell-type dependent activation in the absence of phenotypic skew, we developed an imaging-based method to quantify HA activation at the single-cell and single-virion level. Treating infected airway cultures or viruses with low-pH buffer triggers the transition of activated HA from its pre-fusion to post-fusion conformation, allowing us to quantify activated and unactivated HA using fluorescent prefusion-specific (FI6v3^32^) and post-fusion specific (S1V2-72^50^) antibody fragments (Figure 6A). This approach is analogous to prior work dissecting pH-induced conformational changes in HA using anti-peptide antibodies^51^. To calibrate this approach, we applied it to viral populations with no activation, incomplete activation, or complete activation, as validated by Western blot (Supplementary Figure 11A&B). Following low-pH treatment, we observed a marked increase of the ratio between S1V2-72 and FI6v3 as activation progresses, confirming the method’s sensitivity to conformational changes in HA, even within a compressed range from ∼94% to ∼98% activation resulting from the high efficiency of TPCK-trypsin (Supplementary Figure 11C).

**Figure 6:**
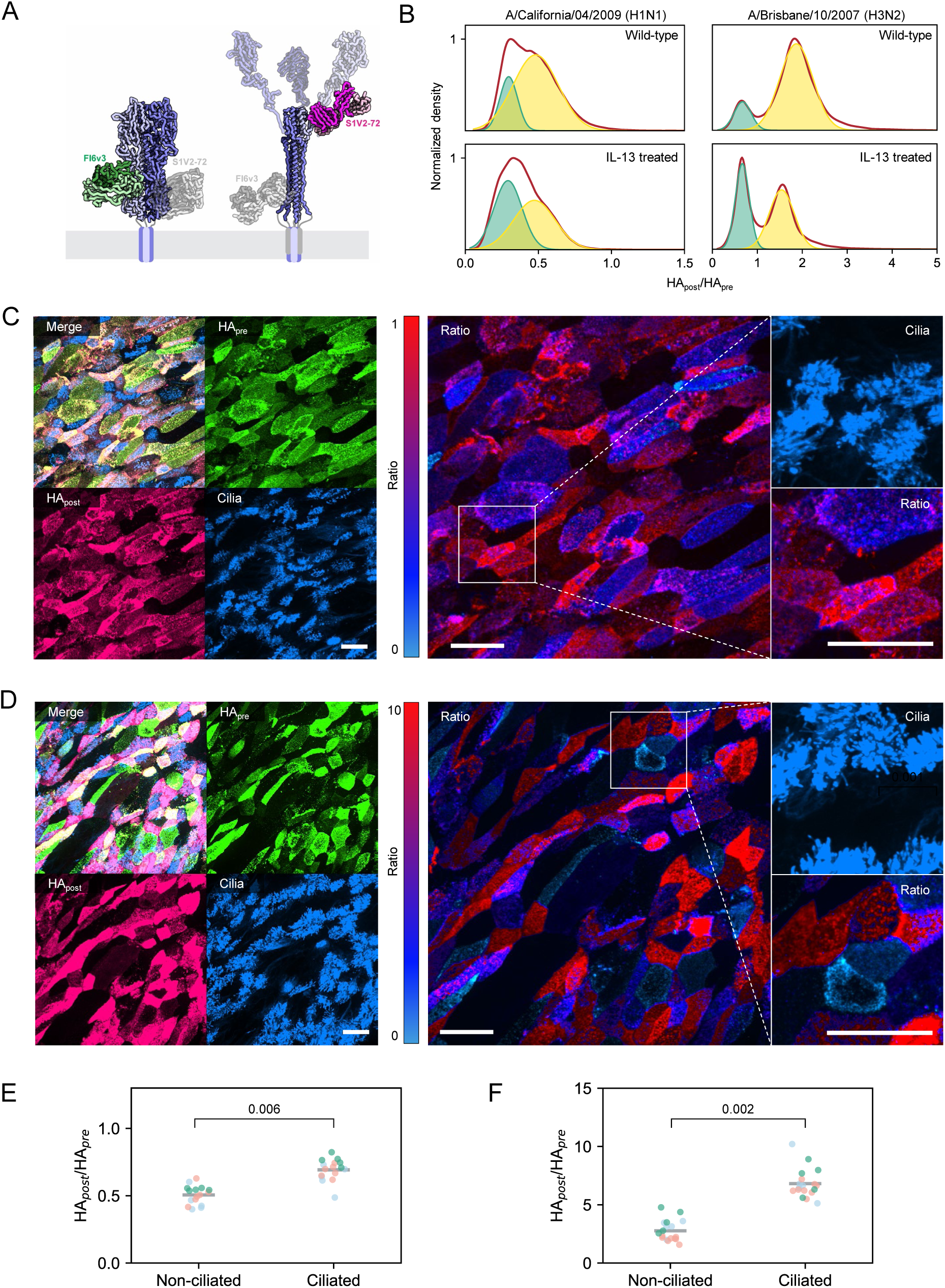
A fluorescence-based approach to measuring HA activation at the single virion and single cell level. (A) Alignments of FI6v3 (PDB ID 3ZTJ) and S1V2-72 (PDB ID 8UDG) binding to their respective epitopes in pre-fusion (left) and post-fusion (right) HA. (B) Distributions of HA_post_/HA_pre_ for individual viruses collected from differentiated HTECs cultured +/−IL-13. Data is fitted with two Gaussian curves. (C) Immunofluorescence and ratiometric (HA_post_/HA_pre_) images of HTECs infected by CA09. Scale bar = 25 μm. Ratio image intensities are modulated by HA channel intensities to highlight infected cells. (D) Immunofluorescence and ratiometric (HA_post_/HA_pre_) images of HTECs infected by BR07. Scale bar = 25 μm. Ratio image intensities are modulated by HA channel intensities to highlight infected cells. (E) Quantification of HA_post_/HA_pre_ on non-ciliated and ciliated cells in differentiated HTECs infected by CA09. Experiments were conducted on three independent samples from donor WU262, depicted by distinct colors in the plot. Each data point corresponds to a single field of view from immunofluorescence confocal imaging. The p-value is determined by a paired t-tests performed on the mean values of the data from all fields of view in each independent experiment. (F) Quantification of HA_post_/HA_pre_ on non-ciliated and ciliated cells in differentiated HTECs infected by BR07. Experiments were conducted on three independent samples from donor WU262, depicted by distinct colors in the plot. Each data point corresponds to a single field of view from immunofluorescence confocal imaging. The p-value is determined by a paired t-test performed on the mean values of the data from all fields of view in each independent experiment.

Next, we analyzed viruses from differentiated HTEC cultures to determine heterogeneity in HA activation. Under standard differentiation conditions favoring ciliogenesis, we identified distinct subpopulations of virions with varying degrees of activation (Figure 6B). This heterogeneity was more pronounced in IL-13 cultures with reduced ciliation. Interestingly, we also observed viruses with high HA_post_/HA_pre_ levels in samples grown from MDCK cells, where no trypsin or Nafamostat is added (Supplementary Figure 12). This aligns with our previous observation that MDCK cells produce a small portion of activated HA that contributes disproportionately to infectious potential (Figure 1E&F).

To investigate the origins of heterogeneity in the activation of virion-associated HA, we applied this imaging approach to differentiated cultures using an anti-acetylated tubulin antibody to distinguish ciliated cells from non-ciliated cells. Ratiometric analysis of HA_post_ and HA_pre_ antibodies revealed a pronounced increase in HA activation on ciliated cells relative to non-ciliated cells following infection with either CA09 (H1N1) (Figure 6C,E & Supplementary Figure 13) or BR07 (H3N2) (Figure 6D,F & Supplementary Figure 13) strains. These findings align with our earlier results using phenotypically skewed cultures and further demonstrate that multi-ciliated cells are not only primary targets of infection for human influenza strains, they are also more effective than other airway epithelial cell types in proteolytically activating HA.

## Discussion

Although proteolytic activation is a critical step in influenza virus replication, the efficiency of activation and its quantitative impact on infectivity in the human airways has not been determined. Using differentiated HTECs as a model, we find that HA activation is efficient but incomplete across the H1N1 and H3N2 strains we tested. Efficient activation is critical for the virus to maintain high infectious potential: our results show a cubic relationship between the level of HA activation and viral infectivity, consistent with a model whereby all three HA monomers within a trimer must be activated to coordinate the conformational changes required for viral entry. Although our results suggest that the potency of protease inhibitors are likely constrained by their ability to access the sites of HA activation, the sensitive dependence of infectivity on the efficiency of HA activation suggests that modest inhibition may be sufficient to restrict multicycle viral replication.

Although previous studies have demonstrated the importance of TMPRSS2 in influenza A virus replication^12,14,24^, the precise determinants of influenza protease dependency remain unclear. One possibility is that protease utilization is encoded within the HA sequence and thus strain- or subtype-dependent. Consistent with this model, an *in vitro* study examining HA activation by trypsin, TMPRSS2, and TMPRSS11D (HAT) revealed that the efficiency of cleavage is dependent on the HA subtype^52^. However, protease dependency does not appear to be directly encoded in the sequence of the HA cleavage site, as prior studies in mice showed that mutations outside this region enable certain H3 strains to utilize alternative proteases for activation^16,41^. An alternate explanation is that specificity may be encoded in additional structural features of the protease and its viral substrates. AlphaFold predictions of unactivated HA and membrane-associated TMPRSS2 show a parallel arrangement of their transmembrane domains, offering a model for how HA could be activated in *cis*. The differences in the extracellular regions of TMPRSS11D or TMPRSS4 may position their serine protease domains in orientations that are less favorable for HA cleavage. Mutations in HA away from the cleavage site could modulate the conformational flexibility of HA to better accommodate alternate proteases. It is worth noting that furin, which cleaves polybasic motifs in highly pathogenic influenza and in SARS-CoV-2, contains a more extensive extracellular region with predicted disorder, which may confer greater flexibility to the serine protease domain as compared to TMPRSS2. Structural and biophysical studies of full-length serine proteases are needed to determine the conformational flexibility of their protease domains and how this might contribute to the breadth of substrates they can accommodate within the structural context of full-length viral fusion proteins.

Finally, we find that HA activation is most efficient within ciliated cells. Consistent with large cell-to-cell differences in HA activation, we do not detect HA-activating proteases shed into apical mucus, which we would expect to make activation more uniform. Previous work has shown disparate outcomes of influenza virus infection in distinct airway cell populations^22,53^. The current work suggests that the efficiency of proteolytic activation is another factor which differs between cell type, with implications for the infectious potential of the viruses that are produced. Interestingly, neither transcriptomic data nor immunofluorescence analysis of protein expression appears predictive of the cell type specific differences in activation we observe. One possibility is that protease activity is subject to complex regulation involving multiple proteins that can either enhance^54^ or suppress^55,56^ HA activation in a cell type dependent manner. Decoding this network could pave the way for antivirals targeting an essential stage in the replication of influenza and other human respiratory viruses.

## Materials and Methods

### Cell lines and viruses

Recombinant viruses were rescued using reverse genetics^57^ as previously discussed. Briefly, virus collected from HEK-293T/ MDCK-II co-cultures transfected with viral plasmids were plaque purified and passaged at MOI ∼ 0.001 in MDCK-II cells in virus growth medium comprised of Opti-MEM (Gibco), 2.5 mg/ml bovine serum albumin (Sigma-Aldrich), 1 µg/ml TPCK-treated trypsin (Thermo Scientific Pierce), and 1× antibiotic-antimycotic (Corning). Infected cells were monitored for cytopathic effect and collected at 24 h.p.i. All virus strains used in this work were sequenced via sanger sequencing for verification purposes.

Cell lines used in this study were purchased as authenticated cell lines (STR profiling) from ATCC and cultured at 37°C with 5% CO_2_ using DMEM (Gibco) supplemented with 10% fetal bovine serum (FBS) (Gibco) and 1× antibiotic-antimycotic.

### Isolation and culture of primary human tracheal epithelial cells

Human tracheobronchial epithelial cells (HTECs) were obtained from excess surgical tissue from lungs donated for transplantation. Tissue use was exempted from regulation as human subject research by the Institutional Review Board of Washington University in Saint Louis. Airway progenitor cells were isolated and cultured as previously described^48^. Briefly, airway epithelial cells were released from airway tissues following incubation in pronase and isolated by differential adhesion. Cells were expanded on plastic culture dishes coated with rat-tail collagen (Corning) and differentiated on collagen-coated transwell supported membranes (Corning) using air-liquid interface (ALI) conditions to generate secretory and multiciliated cells. Cells were treated with 10 ng/mL human recombinant IL-13 (STEMCELL Technologies) starting ALI day 7 for at least 14 days to skew differentiation to MUC5AC-expressing mucous cells.

### Infection of differentiated airway epithelial cells

The apical surface of HTECs grown in ALI culture days were washed with DMEM/F12 and incubated with 250 nM StcE enzyme^27^ for 30 min prior to infection with virus. StcE treatment was not essential for infection, but increased uniformity across experiments. Virus for infections were diluted into 50 μL of DMEM/F12 added to the apical compartment for 1 h at 37°C. Following infection, excess virus was removed and the apical compartment was washed with DMEM/F12. To collect the virus at 20 h.p.i., we added 50 μL DMEM/F12 containing 250 nM StcE enzyme to the apical compartment and incubated with the cells for 50 min before collecting. The StcE enzyme helps the detachment of the virus trapped by tethered and secreted mucins. The media was then gently spun to remove cell debris.

### Immunofluorescence

The apical and basal compartments of airway cultures were quickly rinsed with PBS and fixed using 4% paraformaldehyde (PFA) for 15 min. Fixed cells were permeabilized and labeled overnight at 4°C using the anti-acetylated alpha tubulin antibody 6-11B-1 (mouse IgG, catalog #: sc-23950, Santa Cruz Biotech). Immediately prior to imaging, cells were washed with PBS and the transwell inserts were excised with a razorblade and mounted face-down on a coverslip for imaging with a Ti2 inverted confocal microscope using a 60×, 1.40-NA oil-immersion objective or a 40×, 1.15-NA water-immersion objective.

### Live imaging of differentiated HTECs

Cells were plated and differentiated on the backside of transwell inserts to enable live imaging on an inverted microscope for the soybean trypsin inhibitor binding test. After treating the cells with 1:1000 SiR-Tubulin for 45 min, the insert was placed on a 35 mm glass bottom dish (Cellvis) with a home-designed holder to ensure the cell surface was within the working distance of the 1.15-NA water immersion objective but not touching the glass.

### Ratiometric imaging of infected differentiated HTECs and viruses

To measure HA activation on differentiated HTECs, the apical side of infected cultures were quickly rinsed with PBS at 20 h.p.i. and incubated with citrate-phosphate buffer (pH 5.0) at room temperature for 1 min. After another rinse with PBS, the cells were incubated at 37°C for 60 min with 68 nM FI6v3 scFv (AF488) and 56 nM S1v2-72 Fab (AF555). The cells were then fixed and permeabilized for immunofluorescence staining of cilia.

For ratiometric measurements of virus particles, we first immobilized the virus to 96-well glass bottom plates (Cellvis) coated with biotinylated Erythrina Cristagalli Lectin (Vector Laboratories) as previously described^58^. Each well was then washed with 200 µL citrate-phosphate buffer (pH 5.0) for three times, incubated for 1 min after the last wash for HA activation and washed once with PBS afterwards. Next, the wells were washed three times with 200 µL PBS containing 34 nM FI6v3 scFv (AF488) and 28 nM S1v2-72 Fab (AF555) and moved to the 37°C incubator for 60 min before three additional washes with PBS and fixation with 3.2% PFA for 15 min. After washing with PBS to remove the remaining PFA, the samples were imaged with a Ti2 inverted confocal microscope using a 60×, 1.40-NA oil-immersion objective.

### Immunoprecipitation of native Golgi membranes from differentiated HTEC cultures

HTEC cultures were seeded onto 6.5 mm transwell inserts and cultured under ALI conditions for at least four weeks. Prior to infection, apical compartments of 12 inserts were washed with HBSS to remove mucus and 100 µL of HTEC basic media containing ∼10^5^ PFU of CA09 or BR07 were added to each insert. After incubating for 1h at 37°C and 5% CO_2_, the virus inoculum was removed and the inserts were washed with HBSS and returned to the incubator under ALI conditions. At ∼20 h.p.i., cultures were washed for 1 h with HTEC basic containing StcE and *CpNA* to remove virus particles associated with cells and mucus. Infected cell cultures were washed on ice with PBS before excising each transwell membrane and placing in a Dounce homogenizer containing 2ml of ice cold PBS and Halt protease inhibitor cocktail (Thermo Scientific). Cells and permeable supports were lysed on ice using 20 passes on the homogenizer and the supernatant containing lysed cells (excluding permeable supports) were collected and centrifuged at 1000×g for 5 minutes to pellet nuclei. Cell lysates were mixed with protein A/G magnetic beads that were previously blocked overnight using Intercept (PBS) Blocking Buffer (LI-COR) and incubated with 4A3 anti-GM130 antibody (5 µg antibody per 1 mg of resin). For the ‘non-specific’ condition, beads were coated with an equivalent amount of an RSV-specific antibody in human IgG1 format. After incubating beads and lysates with gentle rotation for 30 min at 4°C, beads were washed quickly three times in PBS, and Golgi membranes were eluted in clean tubes using PBS plus 2% Nonidet P40 substitute (Sigma). Cell membrane fractions were obtained by pelleting the supernatant following depletion of Golgi membranes. Samples were separated on 4-20% SDS PAGE gels and subjected to Western blot analysis.

### SDS PAGE and Western blot

Samples for protein gels were prepared from cell lysates or collected viruses. For virion HA activation analysis, collected virus was heated at 95°C for 10 min in reducing sample buffer before running on a 4-20% SDS-PAGE (Bio-Rad) and transferred onto nitrocellulose membrane (Thermo Scientific). The membrane was blocked with Intercept (PBS) Blocking Buffer and incubated with primary antibodies overnight. The next day, the primary antibodies were removed and the membrane was washed with PBS containing 0.2% Triton-X100 (Sigma) three times before incubating with secondary antibodies for 45 min. After another three washes with PBS plus 0.2% Triton-X100, the membrane was imaged on a LI-COR Odyssey scanning system. Polyclonal Influenza A H1N1 HA Antibody (rabbit IgG, catalog#: PA534929, Thermo Scientific), which binds exclusively to epitopes in HA2, was used to detect CA09 HA. HA Tag Monoclonal Antibody (2-2.2.14) (mouse IgG, catalog#: 26183, Thermo Scientific) was used to detect H3 HA via the HA tag epitope in HA1.

### Recombinant antibodies and PAI-1

VH and VL sequences of antibodies were obtained from the Protein Data Bank and cloned into backbones containing the CH1 (for Fab), CH1-3 (for IgG1), CH2-3 (for VHH-Fc), or CL domain sequences. Heavy chain Fab sequences were modified with a C-terminal ybbR tag for enzymatic labeling^59^ and His_6_-tag for affinity purification. HEK-293T cells at ∼85% confluency were transfected with verified clones and grown in Opti-MEM with 2% FBS (for Fab fragments) or serum-free conditions (for full-length antibodies) for 7 days. Supernatants were collected and purified using Ni-NTA agarose (Thermo Scientific HisPur) for Fab fragments and scFv or A/G agarose (Thermo Scientific Pierce) for full-length antibodies. Eluted antibodies were quantified by UV-Vis, diluted into a labeling buffer (150 mM NaCl, 25 mM HEPES, 5 mM MgCl_2_) and concentrated to ∼1 mg/mL using a Vivaspin 20 centrifugal filter unit (MWCO 10 kDa) (Sartorius). Sfp synthase and CoA-conjugated dyes were prepared as previously described^60^ and used to perform the labeling reaction. Excess dye was removed by PD-10 desalting columns (Cytiva). Recombinant PAI-1 was expressed and purified from HEK293 cells analogous to Fab fragments.

### Recombinant nanobodies, inhibitors and enzymes

A07 nanobody VHH sequences were obtained from the Protein Data Bank and cloned into the pET-28b bacterial expression vector with C-terminal His-tag. BL21 cells were transformed and grown with Kanamycin selection to OD_600_∼0.7. Cultures were then cooled to 25°C and induced with 250 µM IPTG for overnight expression. The protein was purified from the cell lysate using Ni-NTA agarose. Active StcE with an N-terminal His-tag was cloned into pET28b and expressed in BL21 cells similar to the A07 nanobody. Protein was purified using Ni-NTA resin and eluted using 500mM NaCl, 25 mM Tris (pH 7.5), and 250 mM imidazole and stored at 4°C until use.

### Soybean trypsin inhibitor labeling

Soybean trypsin inhibitor was labeled using amine-reactive NHS dye by diluting protein into 100 mM NaHCO_3_ (pH 8.5) buffer and quickly mixing with Janelia Fluor® JF549 SE (Torcris) at a molar ratio of 1:5 (protein:dye). After 10 min incubation at room temperature in the dark, the reaction was quenched by the addition of Tris at a concentration of 10 mM. The labeled protein was then applied to a NAP-5 column (Cytiva) equilibrated with PBS+1mM EDTA for the removal of excess dye.

### Engineered recombinant TMPRSS2 extracellular domain expression and purification

Engineered recombinant TMPRSS2 ectodomain (dasTMPRSS2) was expressed and purified as described previously with some modifications^61^. Briefly, plasmid containing the dasTMPRSS2 ectodomain 109-492 plus EFVEHHHHHHHH (C-terminus) and honey bee mellitin tag (N terminus) was purchased from Addgene (176412). Plasmid was transformed into Escherichia coli D10Bac cells to generate the baculovirus bacmid following the protocol in the Bac-to-Bac® Baculovirus Expression System (Gibco). The resultant bacmid was reversed transfected into Sf9 cells in Sf-900 II SFM (Gibco) using jetOPTIMUS (Polyplus) according to the manufacturer protocol. The generated p0 stock was then passaged sequentially to generate a p1 and p2 stock. Four liters of Sf9 cells at a density of 2.5-4×10^6^ cells/mL were infected with 64 mL of the p2 viral stock. Three days later, the media was collected, buffered with PBS to a final concentration at 1×, pH adjusted to 7.4. The resulting precipitate was removed and the supernatant was applied to 60mL Ni-NTA XPure Agarose Resin (UBPBio). After a two-hour incubation at 16°C with shaking, the resin was applied to a gravity flow column at 4°C, washed and eluted with 1× PBS pH 7.4 containing 500mM imidazole. Fractions containing dasTMPRSS2 were concentrated in an Amicon Ultra centrifugal filter unit (MWCO 30 kDa) (Millipore) and buffer exchanged to less than 5 mM imidazole with SEC buffer (50 mM Tris [pH 7.5], 250 mM NaCl). The protein was left on ice overnight for activation before further purification via a HiLoad 16/600 Superdex 75 pg (Cytiva) size exclusion column at 4°C using SEC buffer. Fractions containing the dasTMPRSS2 were concentrated to 10 mg/mL and stored in SEC buffer with 25% Glycerol at −80°C before using.

### Recombinant dasTMPRSS2 inhibition assay

dasTMPRSS2 was diluted in assay buffer (25 mM Tris-HCl [pH 8.0], 150 mM NaCl, 5 mM CaCl_2_, 0.01% Triton X-100) at a concentration of 3 nM. To calculate K_M_, Boc-QAR-AMC (Bio-Techne) was serially diluted in DMSO such that the final substrate concentration in the assay ranged from 500 μM to 12.5 μM. Assays were performed in a total volume of 30 μL in triplicate wells of black 384-well plates (Nunc), and initial velocity was measured in 35-s intervals for 7 min with excitation of 360 nm and emission of 460 nm on a BiotekHTX plate reader. K_M_ and V_max_ were calculated from Michaelis-Menten plots using GraphPad Prism.

For inhibition assays, inhibitors were serially diluted 4-fold in DMSO, and then further diluted in assay buffer to achieve a final DMSO concentration of 2%. Inhibitor concentrations ranged from 2 μM to 0.15 pM for all tested compounds. The final concentration of substrate was 16 μM, which was the calculated K_M_ value. Assays were performed in a total volume of 40 μL in triplicate wells of black 384-well plates, and initial velocity was measured in 35-s intervals for 7 min with excitation of 360 nm and emission of 460 nm on a Biotek HTX plate reader. IC_50_ values were reported as the average of two independent trials and were calculated from dose-response curves using Graph-Pad Prism.

### Measuring the relationship between HA activation and infectious potential

MDCK cells were infected at an MOI ∼ 1 and cultured in Opti-MEM (Gibco) with 1× antibiotic-antimycotic (Corning) but without TPCK-treated trypsin for 20 hours before collecting the virus-containing media. Detached cells were removed by a spinning at 2000xg for 5 min. The virus was kept at 4°C until ready to use. A day before the virus binding and infection experiments, A549 cells were plated at a confluency of 50% onto 8-chambered cover glass (Cellvis) treated with human plasma fibronectin (Millipore Sigma) according to the manufacturer’s protocol.

To prepare viruses with different HA-activation levels, viruses from trypsin-free cultures were treated with 0.5 µg/mL TPCK-trypsin (Thermo Scientific Pierce) for 5 min, 10 min, 15 min, 20 min, 25 min, 30 min at room temperature. The treatments were terminated by adding soybean trypsin inhibitor (Gibco) at a final concentration of 2.5 µg/mL and mixed immediately. In addition, baseline activation was determined by directly mixing the virus with soybean trypsin inhibitor at 0 min. For fully-activated samples, virus was incubated with 0.5 µg/mL TPCK-trypsin for 45 min at 37°C. HA-activation levels were determined by Western blot using anti-HA antibody PA5-34929 (Invitrogen). The ratio between the intensity of HA2 and HA0, obtained from Image Studio 6.0 software (Licor) was calculated and used as the level of HA activation.

To measure the relationship between HA activation and the efficiency of virus infection, we added serial dilutions of viruses with different HA activation levels to A549 cells and incubated for 30 min at 37°C. Afterwards, the virus-containing media was replaced with fresh Opti-MEM containing 7 nM FI6v3 Fab fragment labeled with Sulfo-Cy5 to block secondary infection and visualize primary infection. At 18 h.p.i., six fields of view were randomly selected on the confocal microscope with a 10×, 0.45-NA objective. Numbers of infected cells were determined by the ratio of HA-positive region over the total area of the field of view.

### Measuring the relationship between virus binding and HA activation

To test whether HA activation affects virus binding, viruses were diluted in Opti-MEM to the same concentration as used in the infection experiment and labeled with 7 nM FI6v3 Fab fragment (Sulfo-Cy5) for visualization 15 min before adding to A549 cells. To visualize cell nuclei, the virus-containing media also contained Hoechst 33342 (Thermo Scientific). Cell growth media was removed, and the cells were washed with PBS before incubating with the viruses. After 30 min of incubation, the virus-containing media was replaced with fresh Opti-MEM after washing with PBS. The number of viruses bound per cell was determined by measuring the numbers of total viruses and total cells per field of view collected with a 40×, 1.30-NA objective on a Nikon Ti2 equipped with a spinning-disk confocal unit. For analysis, maximum-intensity projections for each field of view were used.

### AlphaFold2 modeling of the HA0-TMPRSS2 complex

A structural model of H1 HA in complex with TMPRSS2 was obtained using ColabFold^62^, using the full-length CA09 HA0 sequence as an input along with human TMPRSS2 (80-255) and TMPRSS2 (256-492), reflecting its activated form following cleavage, but omitting the disordered intracellular domain. Models were generated using PDB ID 1HA0^63^ as a custom template, to prevent the model from converging on structures in which the HA cleavage loop is buried within the trimer (as found in structures of processed HA). The top-ranked model after relaxation is shown in Figure 2 and was used for subsequent analysis.

### Single cell analysis

An integrated cell atlas of the human lung in health and disease (core) data^64^, including data of healthy lung tissue from 107 individuals is downloaded from CZ CELLxGENE (https://cellxgene.cziscience.com/collections/6f6d381a-7701-4781-935c-db10d30de293). Genes and cell types of interests were selected and plotted using Python Scanpy.

### Data fitting and statistics

Single virus ratiometric data is fitted to the sum of two gaussians using Python Scipy.Optimize.curve_fit function.

All statistical tests were performed in Python Scipy Stats. No statistical methods were used to predetermine sample size. Statistical tests and the number of replicates used in specific cases are described in figure captions.

## Acknowledgements

This work was supported by NIH R01AI171445.

## Supplementary Figures

**Supplementary Figure 1:**
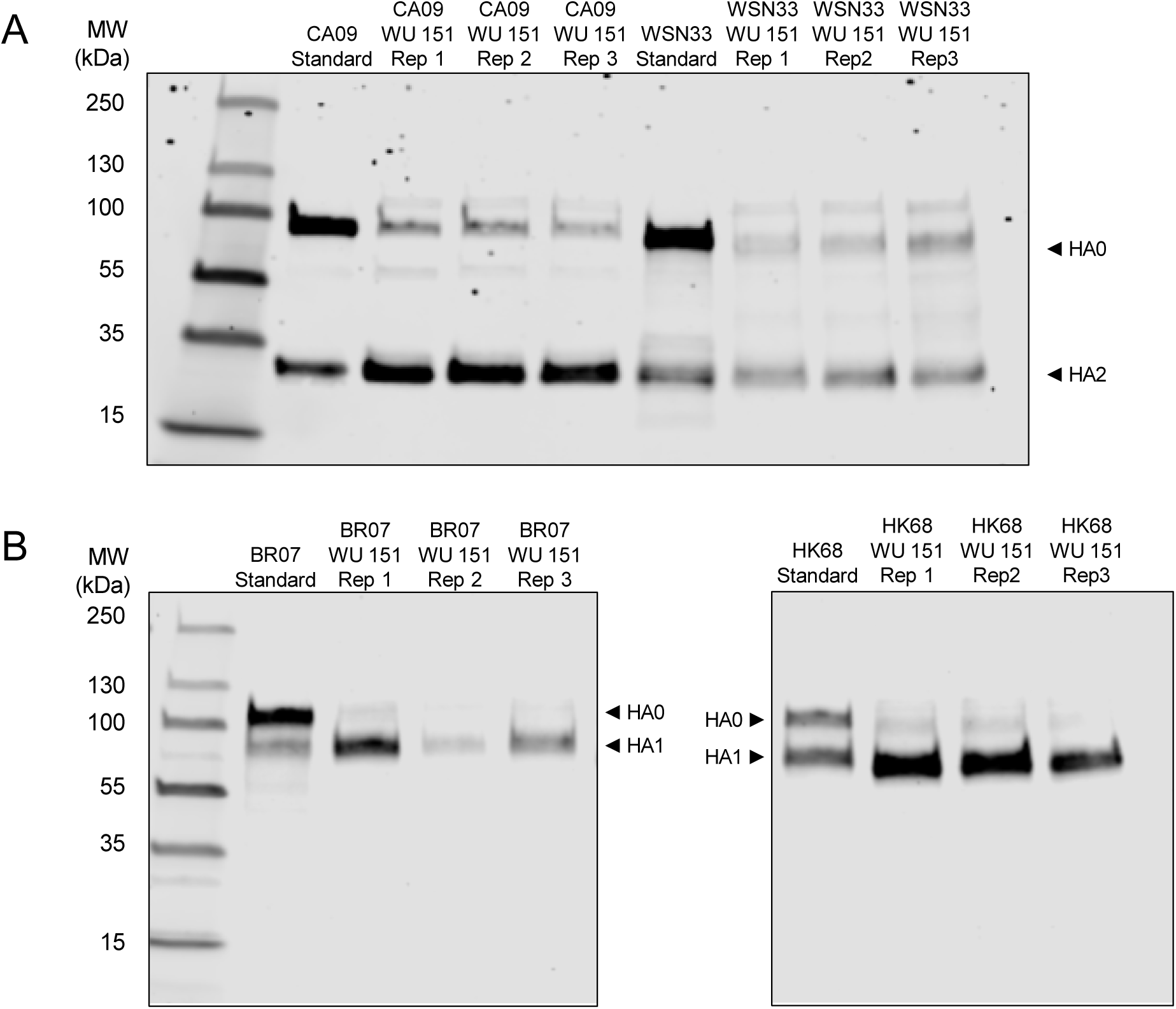
Western blot replicates used to quantify HA activation in differentiated HTEC cultures. All experiments were performed in cells donated by the same patient (WU151). (A) Results for H1 strains (B) Results for H3 strains. Blots are cropped to remove lanes not used in the experiment.

**Supplementary Figure 2:**
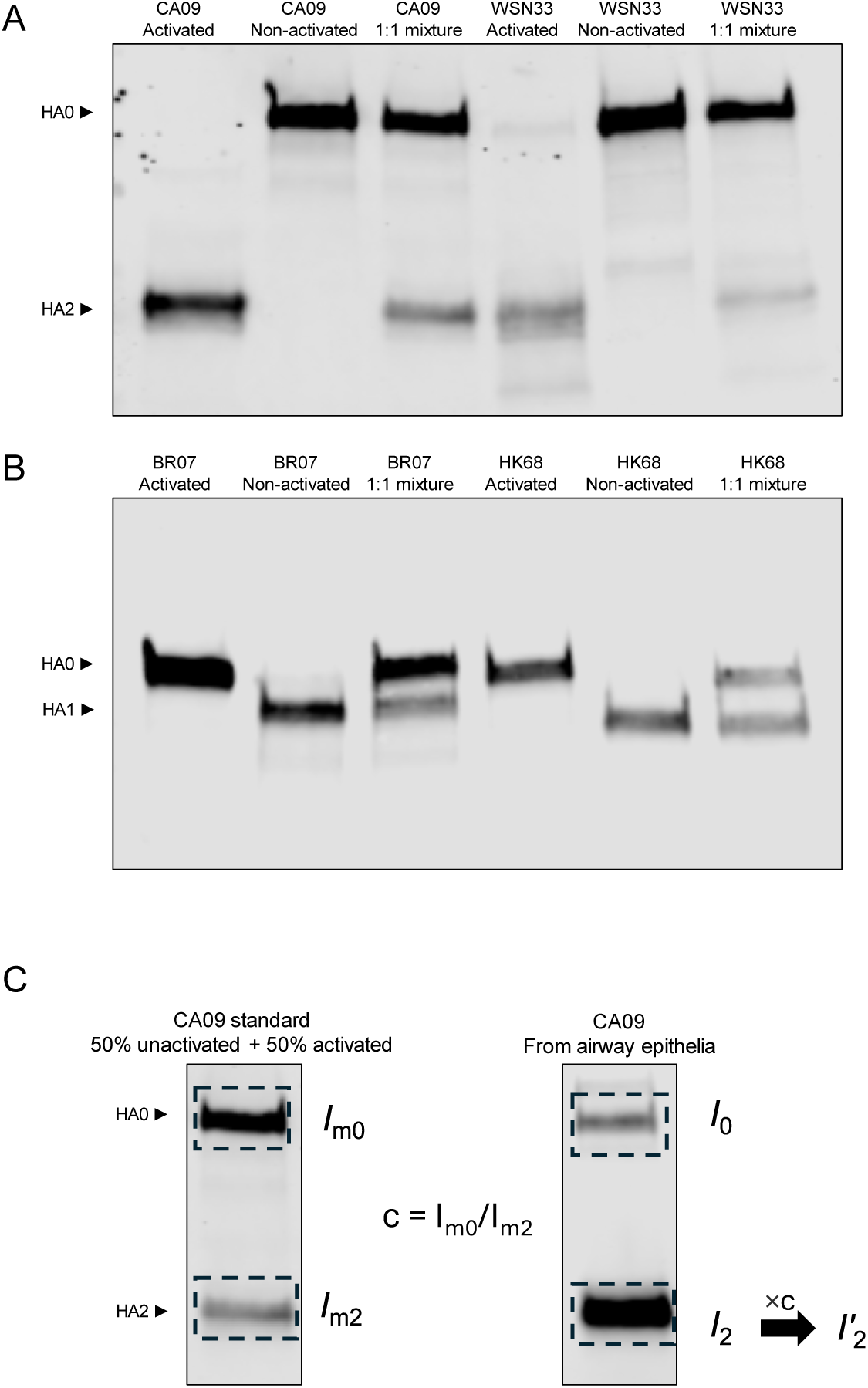
Calibrating Western blot transfer efficiency using samples with equal amounts of unactivated and activated viruses. (A) Western blot comparing fully-activated virus, unactivated virus, and a 1:1 mixture of activated and unactivated virus using H1N1 strains (CA09, WSN33) probed with antibody recognizing an epitope in HA2. (B) Same as in *A*, but for H3N2 strains (BR07, HK68) probed with antibody recognizing an epitope in HA1. (C) Determining the correction factor (‘*c*’) for activated / unactivated band intensities. The activation ratio is determined by the calibrated intensity of the activated band (I’_2_) and the intensity of the non-activated HA band (I_0_).

**Supplementary Figure 3:**
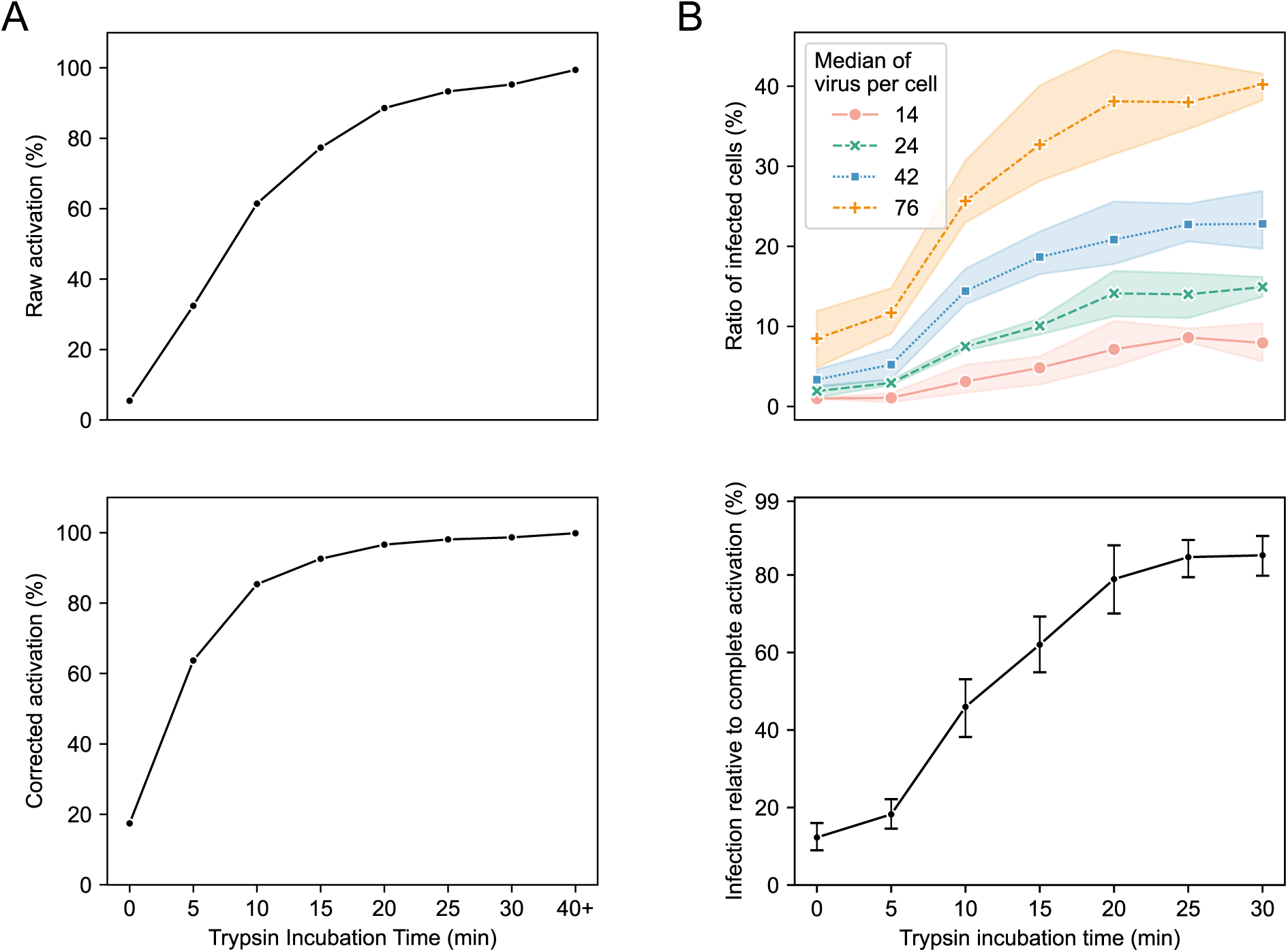
Quantification of HA activation and infectivity of CA09. (A) Quantification of HA activation for CA09 virus incubated with trypsin for different lengths of time. Data is from the Western blot in Figure 1D. Top: uncalibrated raw quantification of HA activation. Bottom: quantification accounting for different detection efficiencies of activated / unactivated bands. (B) Top: Percentage of infected A549 cells following challenge with different amounts of virus, activated by trypsin for different amounts of time. Each curve shows experiments in which different amounts of virus was added and quantified based on the median number of virus per cell. Data is from three replicates. Bottom: Combined data from the top panel showing the relationship between percent infectivity and trypsin incubation time. Percent infectivity is determined relative to infection from samples with complete HA activation.

**Supplementary Figure 4:**
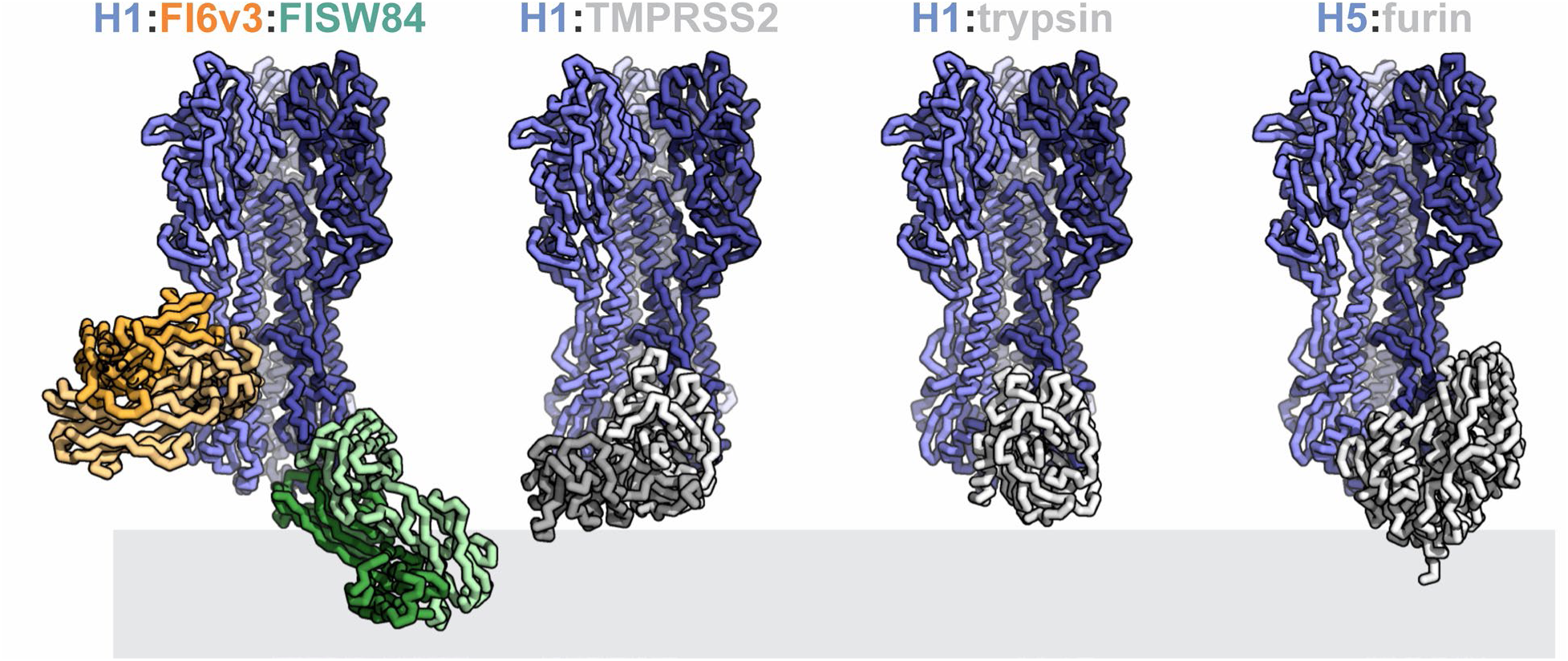
AlphaFold2 models of serine proteases with HA predict steric clashes with stalk- and anchor-binding antibodies. Structures depict the unactivated HA trimer (blue) in complex with one of three proteases (human TMPRSS2, bovine trypsin, or human furin; shown in gray) or FI6v3 and FISW84 Fab fragments.

**Supplementary Figure 5:**
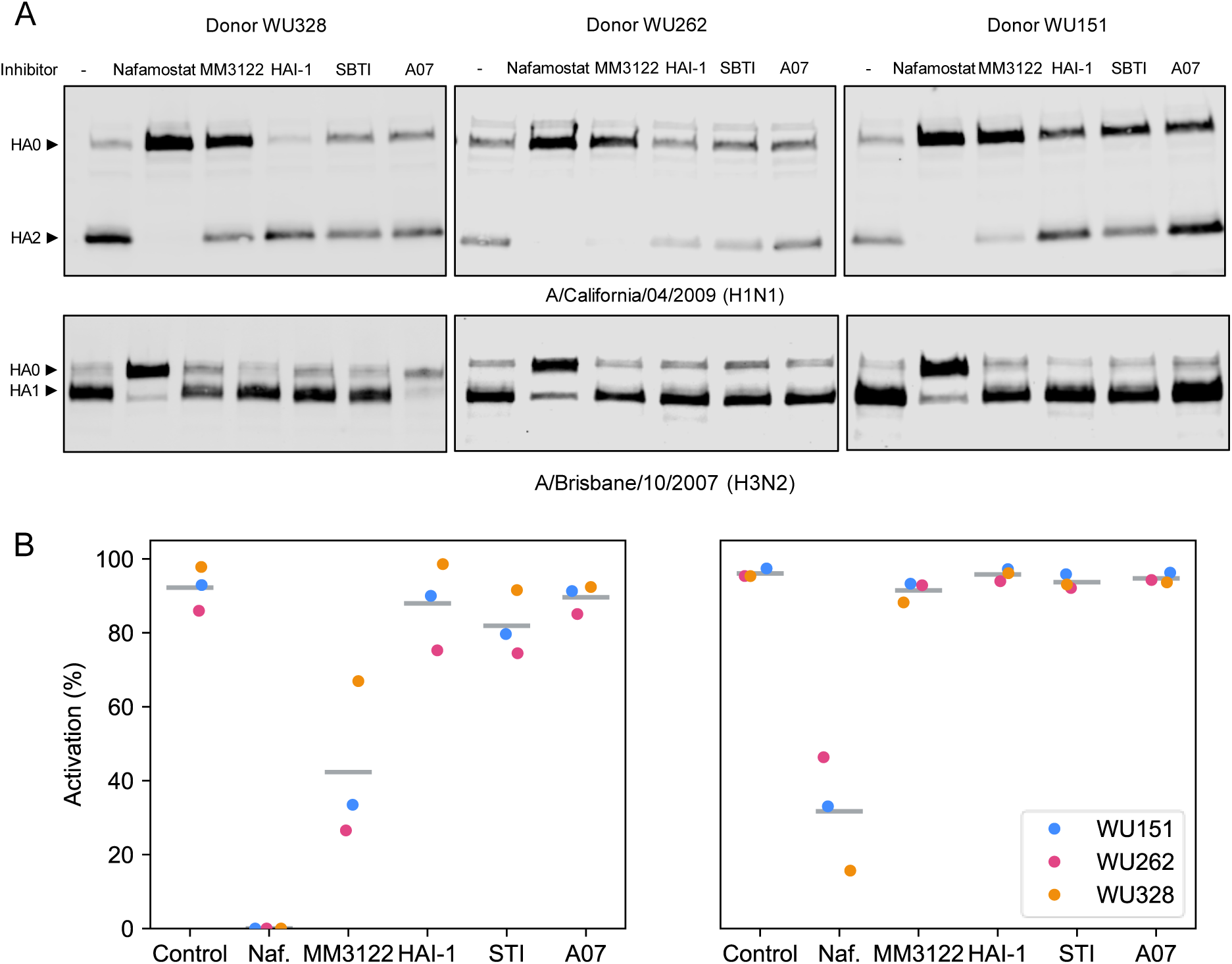
Inhibition of HA activation in differentiated HTEC cultures by selected protease inhibitors. (A) Western blots for quantifying HA activation inhibition of CA09 and BR07 strains by various treatments. Differentiated cultures are derived from three donors. (B) Quantification of Western blots shown in *A*. Values are calibrated based on differences in detection efficiencies between activated / unactivated bands.

**Supplementary Figure 6:**
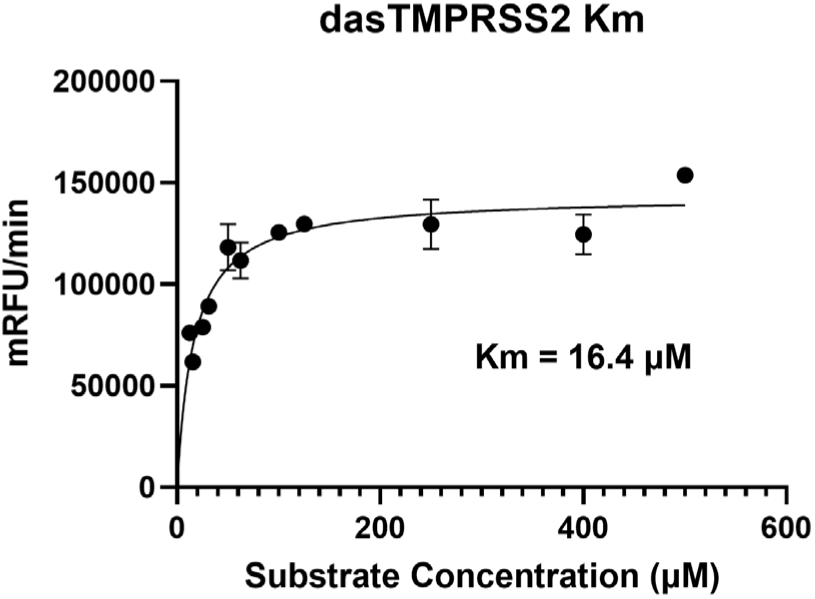
Michaelis-Menten plot of dasTMPRSS2, assayed at a final enzyme concentration of 3 nM.

**Supplementary Figure 7:**
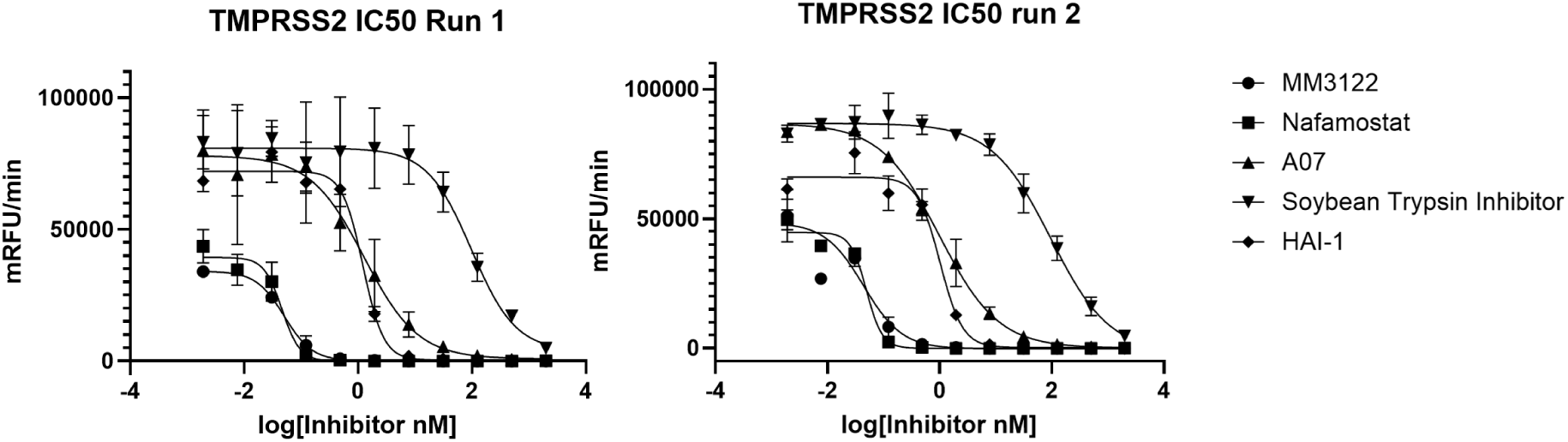
IC50 curves of tested compounds, assayed at a final enzyme concentration of 3 nM and final substrate concertation of 16 μM.

**Supplementary Figure 8:**
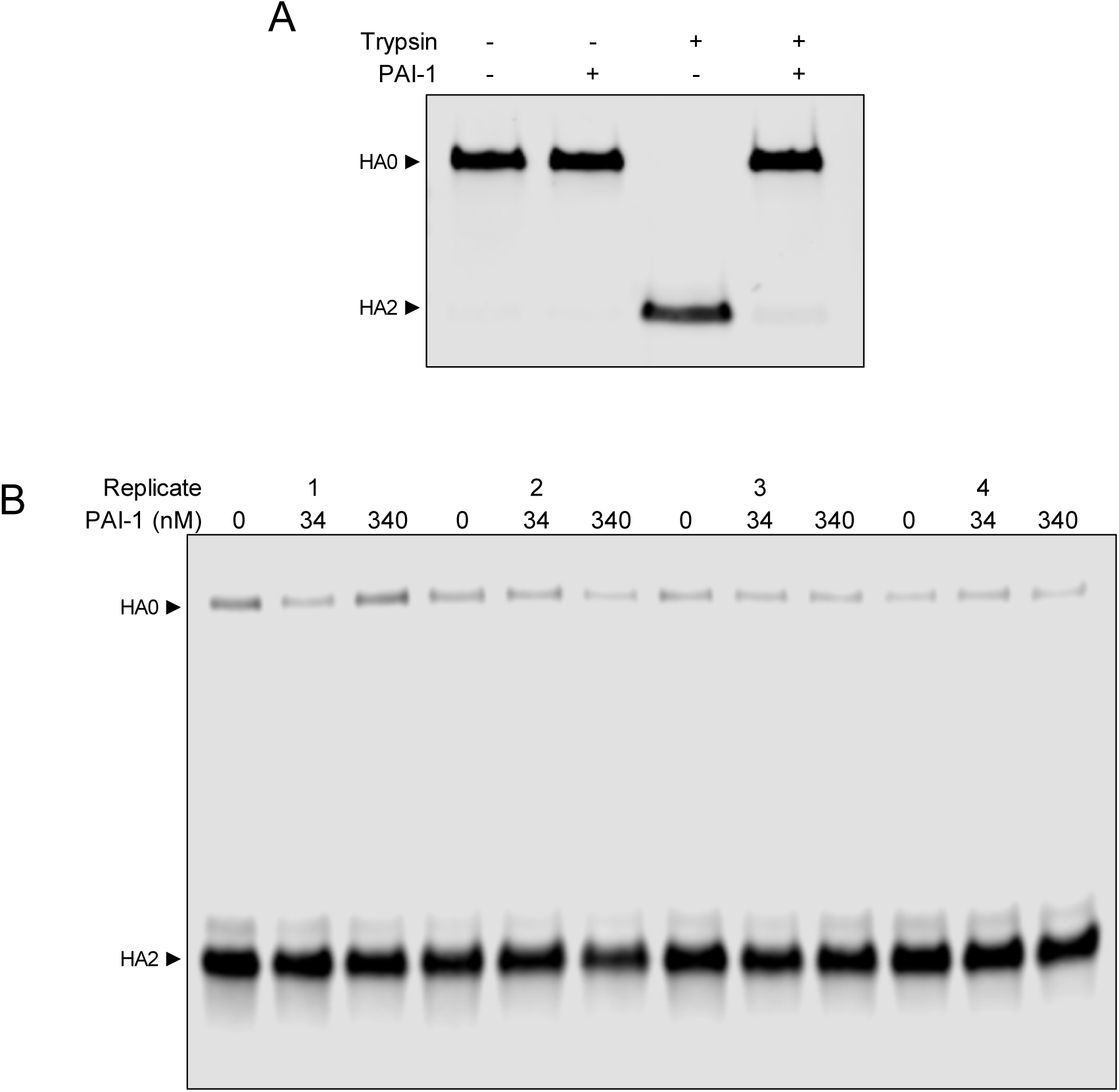
Inhibition of HA activation in differentiated HTEC cultures using PAI-1. (A) Western blot of *in vitro* inhibition tests of 340 nM PAI-1 using CA09 (H1N1). TPCK-trypsin is used at 1 μg/mL (∼40 nM). (B) Western blot of PAI-1 inhibition test in differentiated HTEC cultures using CA09. No inhibition is observed at any concentration tested. Data represent experiments from a total of 12 cultures from a single donor.

**Supplementary Figure 9:**
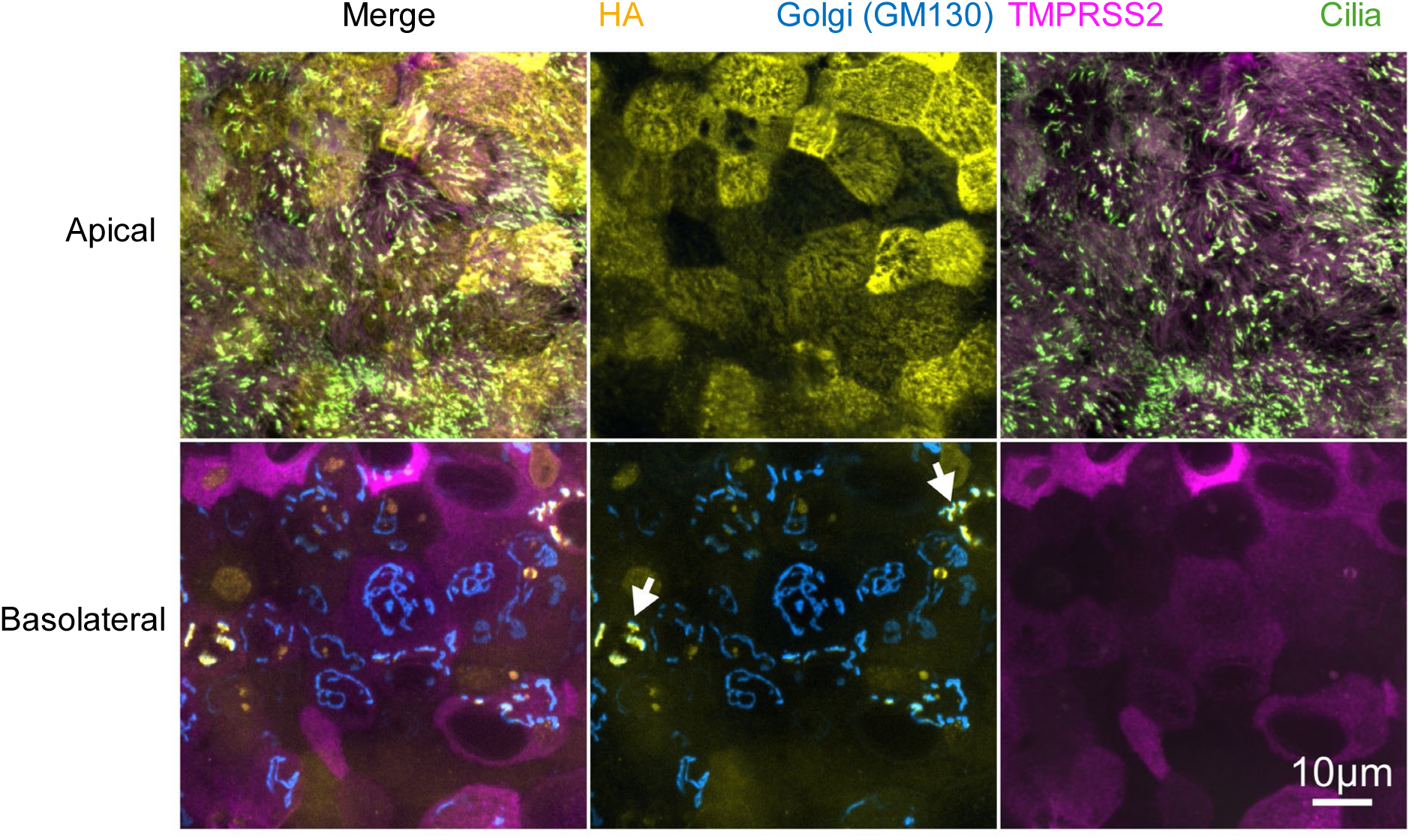
Localization of HA and TMPRSS2 in the Golgi compartment. Immunofluorescence confocal images showing the expression of HA, Golgi marker GM130, and TMPRSS2 in ciliated and non-ciliated cells in a differentiated HTEC culture infected by CA09. Images show apical and basolateral sections of the same field of view.

**Supplementary Figure 10:**
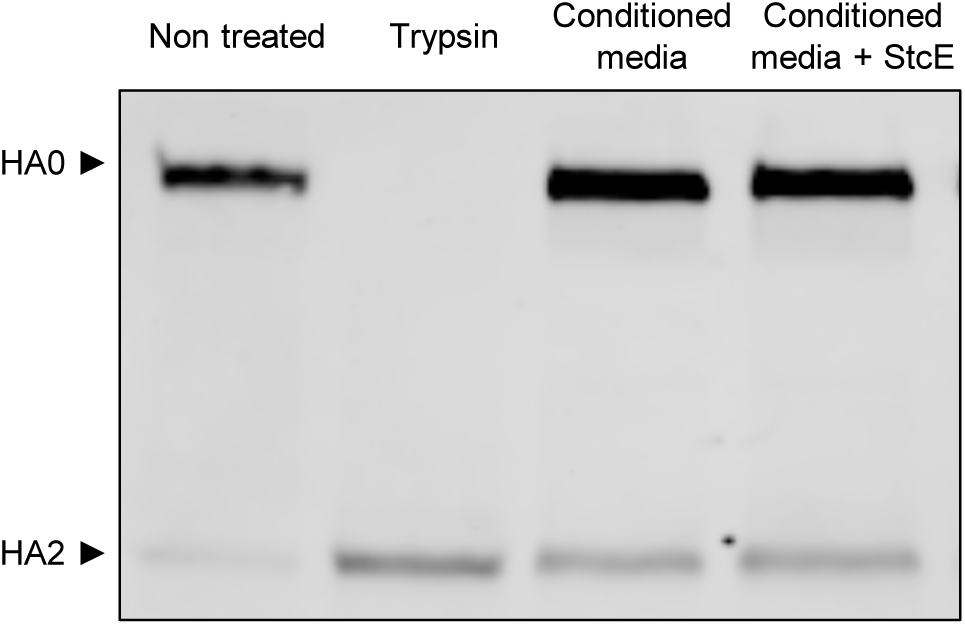
Apical secretions from differentiated HTECs do not efficiently activate HA. Western blot of initially unactivated virus from MDCK cells following control treatment, treatment with trypsin, or treatment with apical secretions from differentiated HTEC cultures (+/− StcE). Experiment is performed with conditioned media from cells of donor WU151.

**Supplementary Figure 11:**
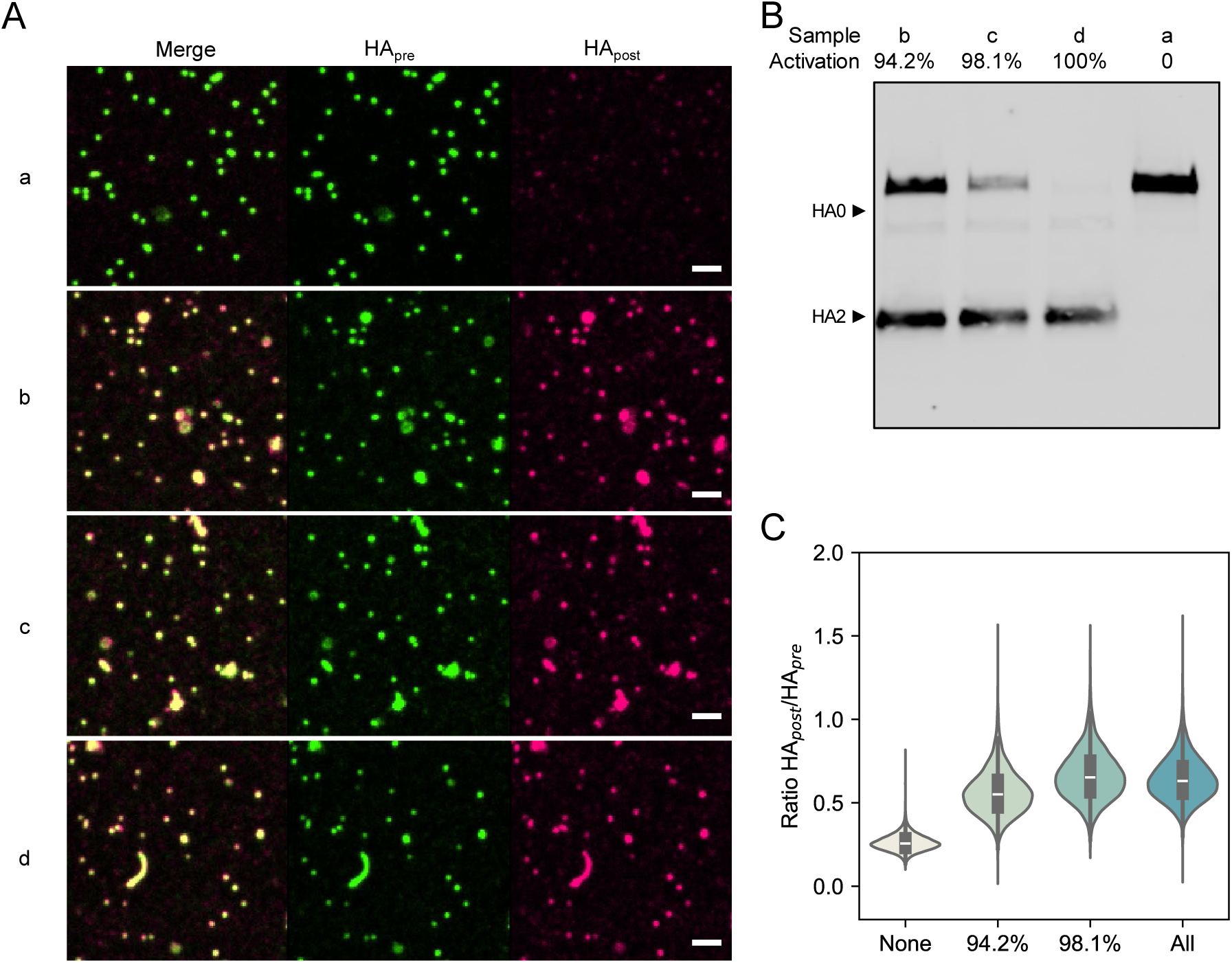
Ratiometric labeling with pre- and post-fusion specific antibodies to quantify HA activation. (A) Representative fluorescence images of viruses activated with trypsin for different amounts of time. Scale bar = 2 μm. (B) Western blot of virus samples shown in *A*. (C) The quantification of the ratio between HA_post_ and HA_pre_ from the fluorescence images. Data are pooled from single virion data from at least five fields containing a total of at least 10^4^ virus particles.

**Supplementary Figure 12:**
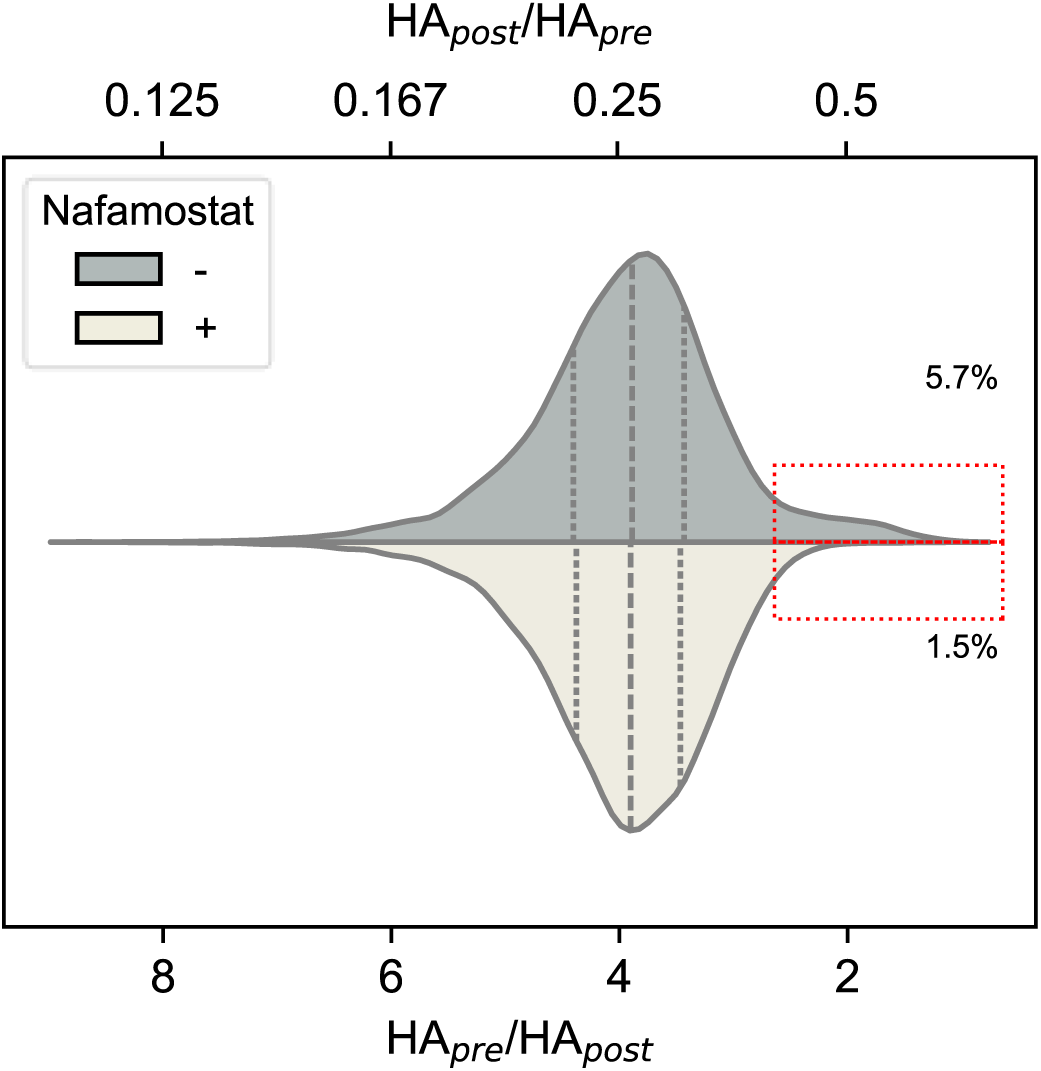
Quantification of baseline HA activation of CA09 virus grown in MDCK cells. A small population of virions (red dashed box) show high activation in the absence of protease inhibitor. This population is suppressed in the presence of 100 nM Nafamostat.

**Supplementary Figure 13:**
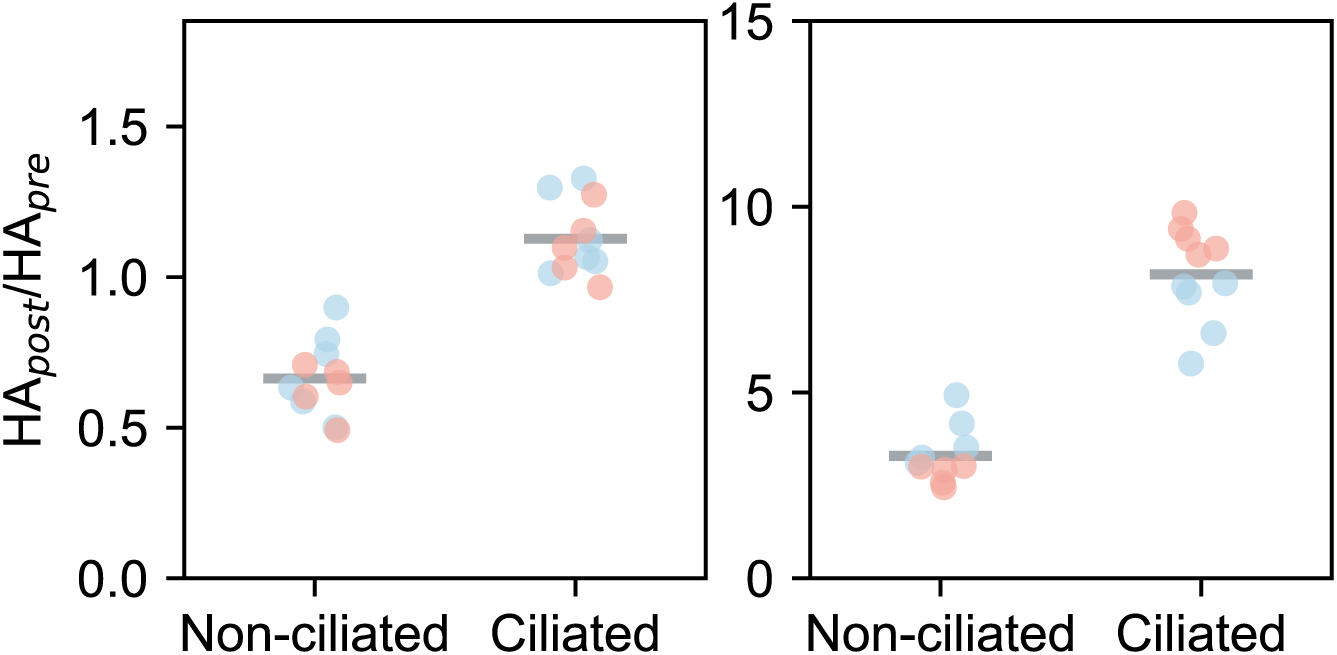
Quantification of HA_pre_/HA_post_ on non-ciliated and ciliated cells in differentiated HTECs infected by CA09 (left) and BR07 (right). Experiments were conducted on two independent samples from donor WU267, depicted by distinct colors in the plots. Each data point corresponds to a single field of view from immunofluorescence confocal imaging. Statistical tests are not available due to the limited number of donor cultures.

